# PopDel identifies medium-size deletions jointly in tens of thousands of genomes

**DOI:** 10.1101/740225

**Authors:** Sebastian Niehus, Hákon Jónsson, Janina Schönberger, Eythór Björnsson, Doruk Beyter, Hannes P. Eggertsson, Patrick Sulem, Kári Stefánsson, Bjarni V. Halldórsson, Birte Kehr

## Abstract

Thousands of genomic structural variants segregate in the human population and can impact phenotypic traits and diseases. Their identification in whole-genome sequence data of large cohorts is a major computational challenge. We describe a novel approach, PopDel, which jointly identifies deletions of about 500 to at least 10,000 bp in length in many genomes together. PopDel scales to tens of thousands of genomes as we demonstrate in evaluations on up to 49,962 genomes. We show that PopDel reliably reports common, rare and *de novo* deletions. On genomes with available high-confidence reference call sets PopDel shows excellent recall and precision. Genotype inheritance patterns in up to 6,794 trios indicate that genotypes predicted by PopDel are more reliable than those of previous SV callers. Furthermore, PopDel’s running time is competitive with the fastest tested previous tools. The demonstrated scalability and accuracy of PopDel enables routine scans for deletions in large-scale sequencing studies.

## Introduction

Comprehensive and reliable collections of genetic variation are a foundation for research on human diversity and disease^1^. They facilitate a wide range of studies investigating mutation rates^2–4^, mutational mechanisms^5–7^, functional consequences of variants^8–10^, ancestry relationships^11^, disease risks^12^, or treatment options^13^. Due to increased throughput and decreased cost, whole-genome sequencing (WGS) is now performed on cohorts of thousands of individuals, for example, sequencing at the population level in Iceland^14^, the United Kingdom^15^, or Crete^16^ as well as sequencing of large cohorts for specific diseases, such as autism^17^ or asthma^18^, and in the general health research context in projects like GnomAD^19^ or TopMed^20^. To create meaningful collections of genetic variation, the data from these large numbers of individuals needs to be integrated. The most direct way of achieving this is done in *joint* variant calling approaches: detecting variant positions and inferring variant genotypes from data of many individuals together.

For single nucleotide variants (SNVs) and small insertions/deletions (indels), joint calling has become the state of the art with tools that scale to tens of thousands of individuals^21,22^. For structural variants (SVs), the analysis of increasingly large numbers of individuals remains a major bioinformatic challenge^23^. Jointly detecting SVs in up to hundreds of individuals is a great achievement of previous projects and tools^24,25^. However, for larger cohorts, catalogues of SVs are generally created in a multi-step approach by first analyzing the data of each individual separately or in small subsets of individuals, subsequently merging the resulting call sets and, finally, determining genotypes for all individuals on the merged call set^26,27^. Merging of SV call sets across individuals is often problematic and arbitrary when the same SV is detected with shifted positions in several individuals^28,29^. In addition, variants that are only weakly supported by the data may not be discovered using this multi-step approach. Furthermore, the aligned read data is accessed at least twice in the process, for detecting and for genotyping SVs, requiring substantial computational resources. A joint SV detection approach simplifies the calling process, is computationally more efficient if accessing the large amounts of input data only once, eliminates the need for an error-prone variant merging step, and may reveal weakly supported variants if carried by several individuals as the support can be accumulated across individuals.

To overcome the limitations of current SV callers, we introduce a joint calling approach for deletions of a few hundred up to tens of thousands of base pairs (bp) in length. The approach is specifically designed for large cohorts and is, to our knowledge, the first joint approach for SV discovery that scales to tens of thousands of individuals. Nevertheless, it can also be applied to a single genome or small numbers of genomes, where it achieves comparable or even better accuracy to widely-used deletion callers.

## Results

Our main result is a novel computational approach that can detect and genotype deletions in tens of thousands of genomes jointly. We have implemented the approach in the tool PopDel and evaluate PopDel’s performance in comparison to previous, popular tools for SV calling on simulated data, publicly available data and population-scale data from Iceland.

### Computational approach for joint deletion calling

Deletions can manifest themselves in the reference alignment of short-read sequences as local drops in read depth, aberrations in the distance between the alignment of two reads in a pair, and split-aligned reads^30^. To detect deletions, PopDel focuses on local changes in the read pair alignment distance compared to the genome-wide distribution of read pair alignment distances. Split-aligned reads are used to infer a precise start position of detected deletions and read depth is used implicitly during genotyping.

To achieve scalability to very large numbers of individuals, PopDel has two steps: A profiling step, which reduces the aligned input sequencing read data per individual into a small read pair profile, followed by a joint calling step, which takes as input the read pair profiles and outputs deletion calls with genotypes across all individuals (Figure 1, Methods). While the two individual steps are novel, the enclosing two-step design is reminiscent of joint calling of small variants in the GATK HaplotypeCaller^21^ and CNV calling approaches that are based on read depth profiles^32,33^. Read pair profiles contain the sequencing experiment’s distribution of read pair distances as well as alignment start positions and distances of all read pairs that match certain quality criteria (**Supplementary Table 1**). The joint calling step processes the profiles of all individuals together in small genomic windows (default 30 bp) to discover and genotype deletions. For all windows, likelihood ratio tests are performed to test if deletions overlap the window in any of the jointly analyzed individuals. In the likelihood computation we use weighted genotype likelihoods to ensure that rare deletions can be found by boosting the signal in carriers and down-weighting the contribution of non-carriers dependent on the allele frequency. Finally, adjacent windows that support the same deletion are aggregated and output together with genotype likelihoods of all individuals.

**Figure 1.**
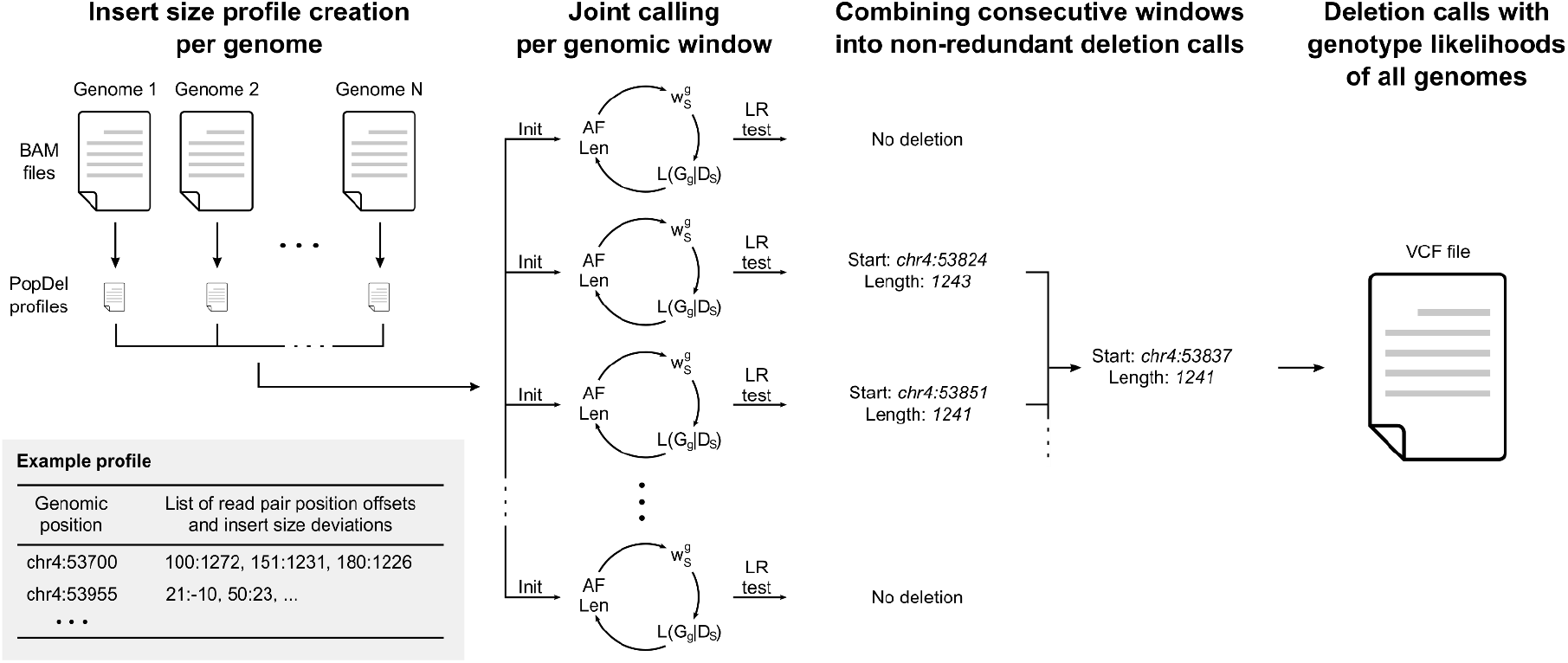
Overview of the approach implemented in PopDel. First, the BAM file of one individual at a time is reduced into a small profile. The profiles of all individuals are processed together by sliding a window (of size 30 bp by default) over the genome and assessing the likelihood of each window to overlap a deletion in any individual. Sizes and allele frequencies of the deletions are estimated iteratively. Consecutive windows are combined into a single variant call and genotypes of all individuals are output to a VCF file. Init, Initialization; AF, allele frequency; Len, deletion length; 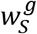, genotype weights per genome; *L*(*G_g_|D_s_*), genotype likelihoods per genome; LR, likelihood ratio.

Most parameters of PopDel are calculated from the input data. The input parameters for each likelihood ratio test are iteratively estimated (Figure 1, Methods): the deletion length, allele frequency, genotype weights and genotype likelihoods for all individuals for the three genotypes (non-carrier, heterozygous carrier and homozygous carrier). The minimum length of deletions that can be identified with our likelihood ratio test derives from the standard deviation of the read pair distances.

### Assessment of scalability on simulated data

For an initial assessment of PopDel’s precision and recall, we simulated two cohorts of sequencing data: the first consisting of 1,000 individuals carrying random deletions and the second consisting of 500 individuals with deletions reported in the 1000 Genomes Project (Methods). Individuals in the first cohort carry on average 659 heterozygous and 673 homozygous deletions that are placed densely on chromosome 21, while individuals in the second cohort carry on average 167 heterozygous and 64 homozygous deletions that are more sparsely distributed on chromosomes 17 to 22. On these data, we compared the recall, precision, running time and memory consumption of PopDel to that of four popular SV callers that can identify SVs jointly in a limited number of individuals or provide a pipeline for single-genome calling followed by merging and genotyping (Delly^34^, Lumpy^35^ via the recommended Smoove pipeline (see URLs), Manta^36^ and GRIDSS^37^). We note that PopDel currently only reports deletions while other callers also report other types of SVs.

The precision and recall of PopDel and most other tools is high for both cohorts (Figure 2) reflecting that the simulated data is easy to analyze. Only GRIDSS’ performance in precision drops significantly with increasing numbers of individuals, which is why we excluded it from further joint calling comparisons. The recall and precision on the 1000 Genomes Project deletions fluctuates more due to the smaller number of simulated deletions per individual. Nevertheless, all tools show a clear trend of higher recall with increasing numbers of individuals on the 1000 Genomes Project deletions. This suggests that these deletions tend to be more difficult to identify than random deletions and, here, deletion callers benefit from integrating data of several individuals. Surprisingly, the recall of Lumpy is higher for the 1000 Genomes Project deletions than for the randomly generated deletions.

**Figure 2.**
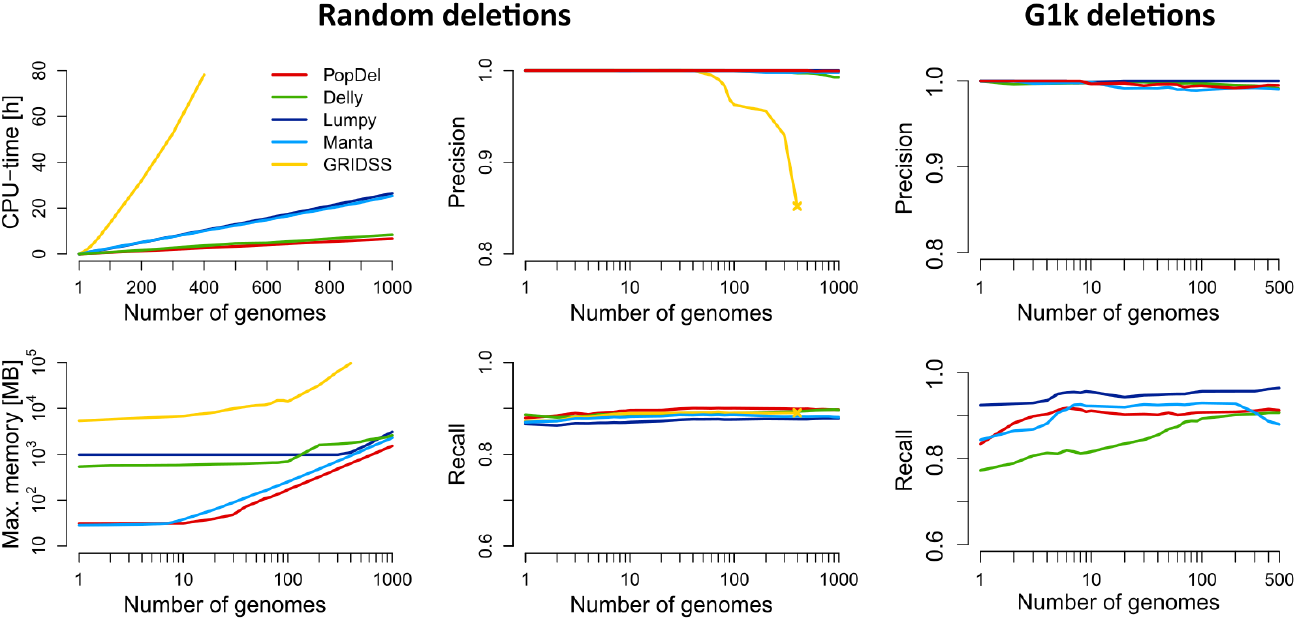
Performance of PopDel in comparison to Delly, Lumpy (via Smoove), Manta, and GRIDSS on simulated data for increasing numbers of genomes. Running time and memory on data with deletions randomly placed on human chromosome 21 (left). Recall and precision on data with random deletions (middle). Recall and precision with deletions reported by the 1000 genomes project simulated on chromosomes 17 to 22 (right).

PopDel is the only tool that we could run on all 1000 individuals using our system’s default settings. For all other tools, we had to increase the ulimit (maximum number of open file handles) in order to complete the calling on more than 500 individuals jointly, indicating that the other tools were primarily designed to jointly analyze smaller numbers of individuals. With a running time of 397 minutes and a peak memory of 1.5 GB for profiling and joint calling on the first cohort of all 1,000 individuals, PopDel is the fastest tool and among the tools that require the least memory (Figure 2).

### Comparison to reference deletion sets from the Genome in a Bottle (GiaB) consortium

Next, we assessed PopDel’s performance compared to Delly, Lumpy (via Smoove), and Manta on short-read WGS data of the well-studied HG001/NA12878 and HG002/NA24385 genomes and their parent’s genomes (Methods). For this assessment we used reference sets of deletion calls prepared by the GiaB Consortium^38^: the short read based reference set and a set of deletions called from PacBio long read data for HG001, and the preliminary variant set for HG002 (see URLs).

PopDel is competitive with the three other tools on the data of HG001 and HG002 (Figure 3, **Supplementary Figure 3 and 4**). All tools succeed to identify the majority of deletions reported in the short read HG001 reference set (661/778, 85.0%) with PopDel identifying marginally more deletions (713, 91.6%) than Lumpy (705, 90.6%), Delly (702, 90.2%) and Manta (680, 87.4%). The fraction of PacBio deletions identified by all three tools is much lower (731/3,831, 19.1%). This is expected as the long PacBio reads reveal variants involving repeats that are invisible or hard to detect in short read data. PopDel identifies a similar number of PacBio deletions (847, 22.1%) as Lumpy (838, 21.9%), Delly (871, 22.7%), and Manta (786, 20.5%). Including those deletions that are not part of the two HG001 reference sets, PopDel reports fewer deletions than Delly, a similar number of deletions as Lumpy, and more deletions than Manta. As the NA12878 reference sets do not claim to be complete, the additional deletions can either be true or false positives.

**Figure 3.**
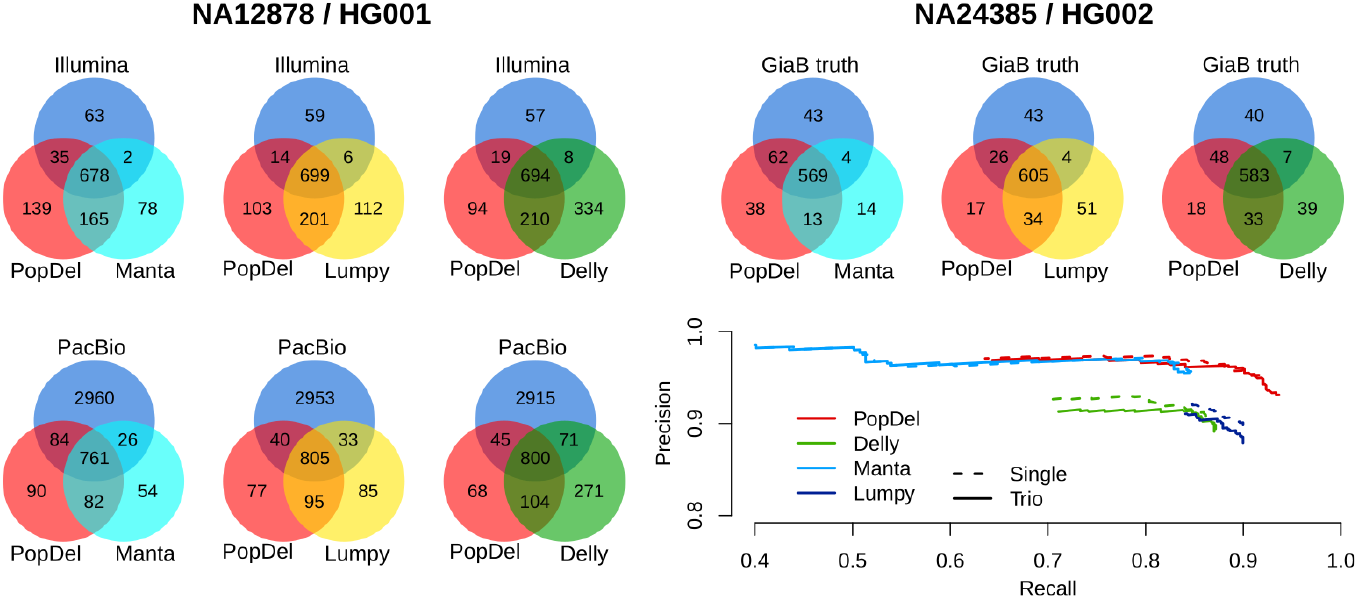
Comparison of PopDel’s accuracy on the Genome-in-a-Bottle benchmark genomes HG001 and HG002 to Manta, Lumpy (via Smoove), and Delly using a minimum reciprocal overlap criterium of 50%. Call set overlap for HG001 with the Genome in a Bottle short read reference set of deletions (top left) and a deletion set called from PacBio long read data (bottom left). Call set overlap for HG002 with deletions from the Genome-in-a-Bottle preliminary variant set (top right). Precision-recall curve when calling deletions from HG002 alone (Single) or jointly with the parental genomes (Trio) for different genotype quality thresholds (bottom right).

The preliminary HG002 deletion set has been released as the first reference set that is near complete within defined high-confidence regions of the genome and, hence, allows us to evaluate the precision and recall of PopDel compared to the other tools (Figure 3). The call set of PopDel, when run on the data of HG002 and its parental genomes jointly, comprises 629 of the 678 reference deletions (93.7% recall) with a precision of 93.1% resulting in an F1 score of 93.4%. Only Manta’s precision is higher (95.5%) at the cost of a much lower recall (84.5%) resulting in an F1 score of only 89.7%, which is similar to that of Delly (88.1%) and Lumpy (88.9%). Thus, PopDel outperforms the other tools by 3.7 to 5.3 percentage points in terms of F1 score. PopDel’s recall is higher when adding the parental genomes compared to running it on the data of HG002 alone (91.6%), indicating a benefit of joint calling. While Manta’s performance is hardly affected by adding the parental genomes to the calling, Delly and Lumpy lose precision on our data without the recall being affected.

### Analysis of population structure based on deletions in the Polaris Diversity cohort

We applied PopDel to the 150 genomes in the Polaris HiSeq X Diversity cohort (BioProject accession PRJEB20654) to evaluate if the deletions identified by PopDel reflect population structure (Methods). The cohort consists of three continental groups: Africans, East Asians and Europeans. PopDel identifies an average of 969 heterozygous and 340 homozygous deletions per individual overall. Consistent with previous studies, Africans carry significantly (P value < 2.2 · 10^−16^, two-sided t-test) more deletions than Europeans and East Asians (Figure 4A). Principal component analysis of PopDel’s deletion calls shows a clear separation between the three continental groups (Figure 4B) mirroring the well-known clustering resulting from small variants^24,39^. In particular, the first principal component separates the African genomes from the other continental groups, while the second principal component additionally pulls apart the European and East Asian genomes. These findings indicate that the deletions detected and genotyped by PopDel well reflect the biological differences between the continental groups. Similar results were obtained for Delly and Lumpy via Smoove (**Supplementary Figure 5**).

**Figure 4.**
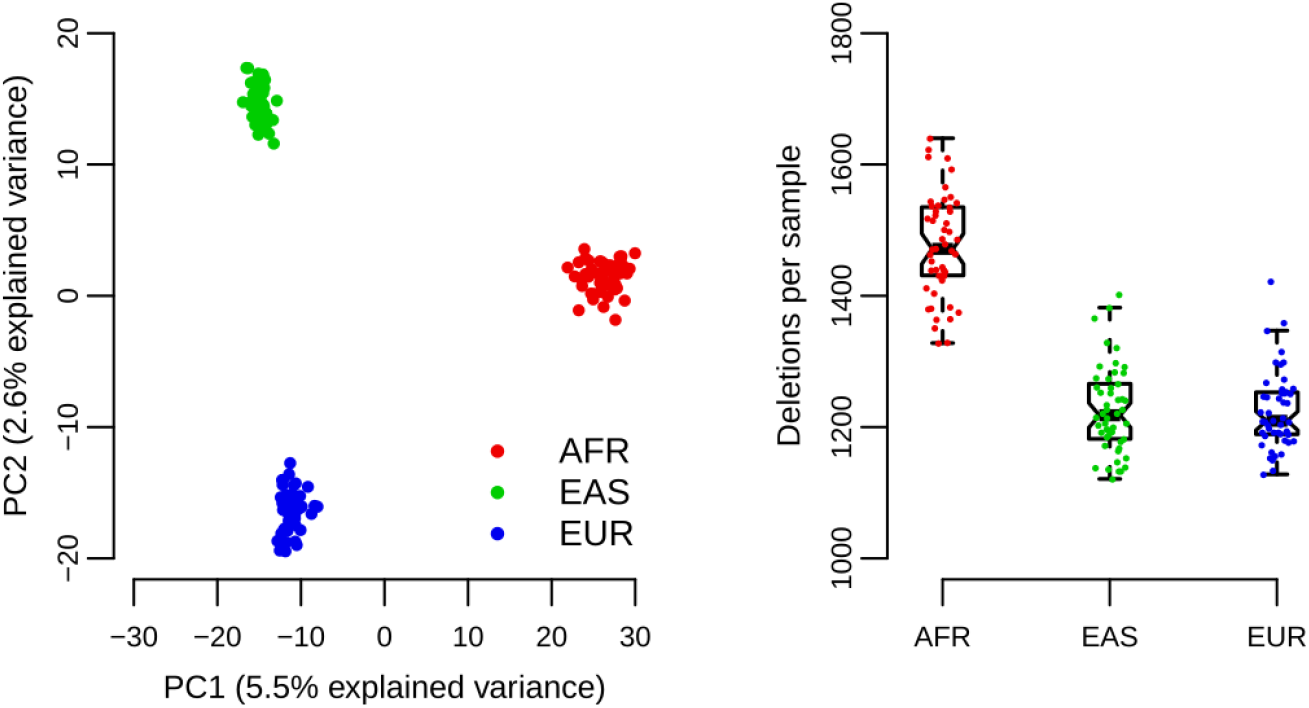
Analysis of PopDel’s deletion calls on the Polaris Diversity cohort consisting of 50 individuals from three continental groups each: principal component analysis (left) and number of deletions per genome (right). AFR, African; EAS, East Asian; EUR, European.

### Evaluation of genotyping using data of 49 Polaris trios

By combining the Polaris Diversity cohort with the Polaris HiSeq X Kids cohort (BioProject accession PRJEB25009) we obtain a set of 49 trios that allow a thorough evaluation of the genotype predictions. After running PopDel, Delly and Lumpy via Smoove (Methods), we analyzed inheritance patterns of deletions and their genotypes in the 49 trios. In particular, we calculated the Mendelian inheritance error rate and transmission rates for each tool (Methods). The Mendelian inheritance error rate effectively assesses the genotyping of common variants. The transmission rate is also meaningful for rare variants measuring how often a deletion allele is inherited from a heterozygous parent (**Supplementary Table 3**), in particular when it is calculated for deletions unique to one trio and the second parent being a non-carrier. As we noted an overabundance of heterozygous deletions in all call sets, we removed deletions that are not in Hardy-Weinberg equilibrium (P value 0.01) before all other calculations.

The genotyping of PopDel is very well calibrated in our comparison to Delly and Lumpy (Figure 5). The deletions called by all three tools can be filtered to Mendelian inheritance error rates below 0.1% using reported genotype quality values. Notably, PopDel reports a larger number of deletions consistent with Mendelian inheritance than Delly and Lumpy when filtering to any Mendelian inheritance error rate. For example, PopDel reports an average of 1177 consistent deletions per trio compared to 1161 (Delly) and 1128 (Lumpy) when filtering to an error rate just below 0.3%. Similarly, PopDel reports more deletions unique to one trio than Delly when filtering by genotype quality to any given transmission rate, for example, 3167 PopDel deletions compared to 2935 Delly deletions when filtering to the best transmission rate closest to 50%. Furthermore, PopDel has a lower Mendelian inheritance error rate than Delly when filtering to the expected transmission rate of 50%. Lumpy’s transmission rate shows a pattern that may be indicative of false positives (**Supplementary Text**) and never reaches 50%.

**Figure 5.**
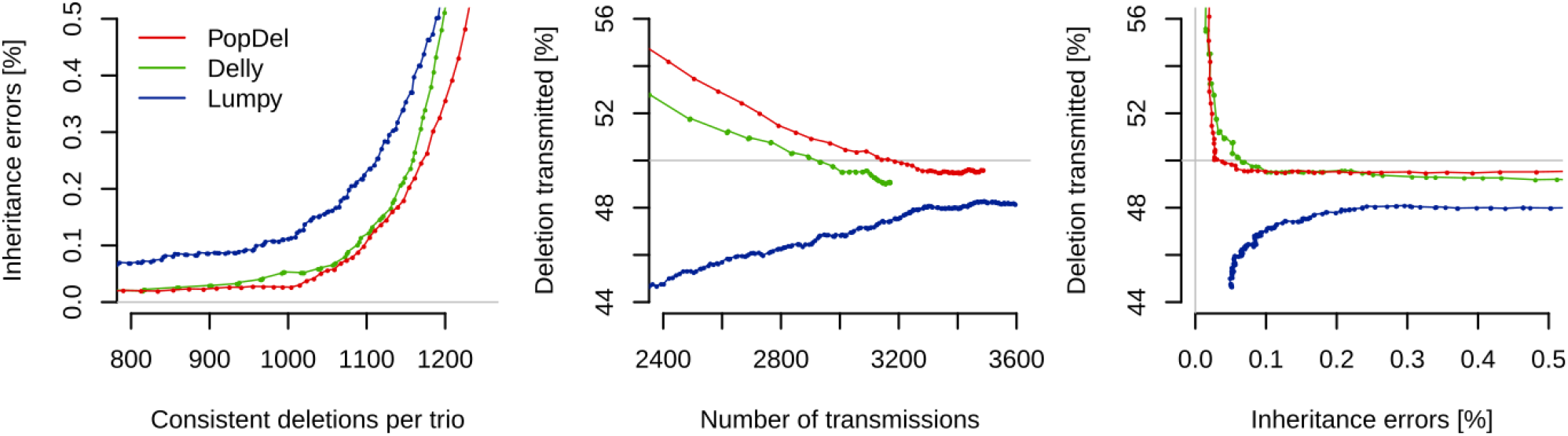
Assessment of genotype inheritance patterns in the Polaris Kids cohort after filtering for Hardy-Weinberg equilibrium (P value 0.01) and varying genotype quality thresholds. Mendelian inheritance error rate by number of deletion sites per trio that are consistent with Mendelian inheritance (left). Percentage of transmitted deletion alleles for all deletions unique to one trio and one parent by the number of possible transmissions, i.e. the number of deletions unique to one trio and one parent (middle). Percentage of transmitted deletion alleles by Mendelian inheritance error rate (right). Gray lines indicate ideal values of the Mendelian inheritance error rate and transmission rate.

We further assessed the genotyping performance based on the rate of transmission from parents to children including scenarios where more than one parent is heterozygous and the deletion was found in several trios. For each tool, we chose genotype quality thresholds that filter the deletions to a Mendelian inheritance error rate just below 0.3% (Table 1). Using these filters, the rate of deletions transmitted from parent to child that are private to a single trio is not significantly different from 50% for PopDel and Delly (two-sided binomial test, P value threshold 0.05). When considering deletions found in several trios where only one parent is heterozygous, the transmission rate of PopDel is not significantly different from the expected 50% (two-sided binomial test, P value threshold 0.05) whereas it is different for Delly when one parent is a homozygous carrier. When both parents are heterozygous, all three tools show an overabundance of heterozygous calls in the child, indicating that this is the most challenging configuration for genotyping. The transmission rate of Lumpy is significantly different from the expected value for all considered genotype configurations. This analysis suggests that PopDel’s genotyping accuracy is superior to that of Delly and Lumpy.

**Table 1.**
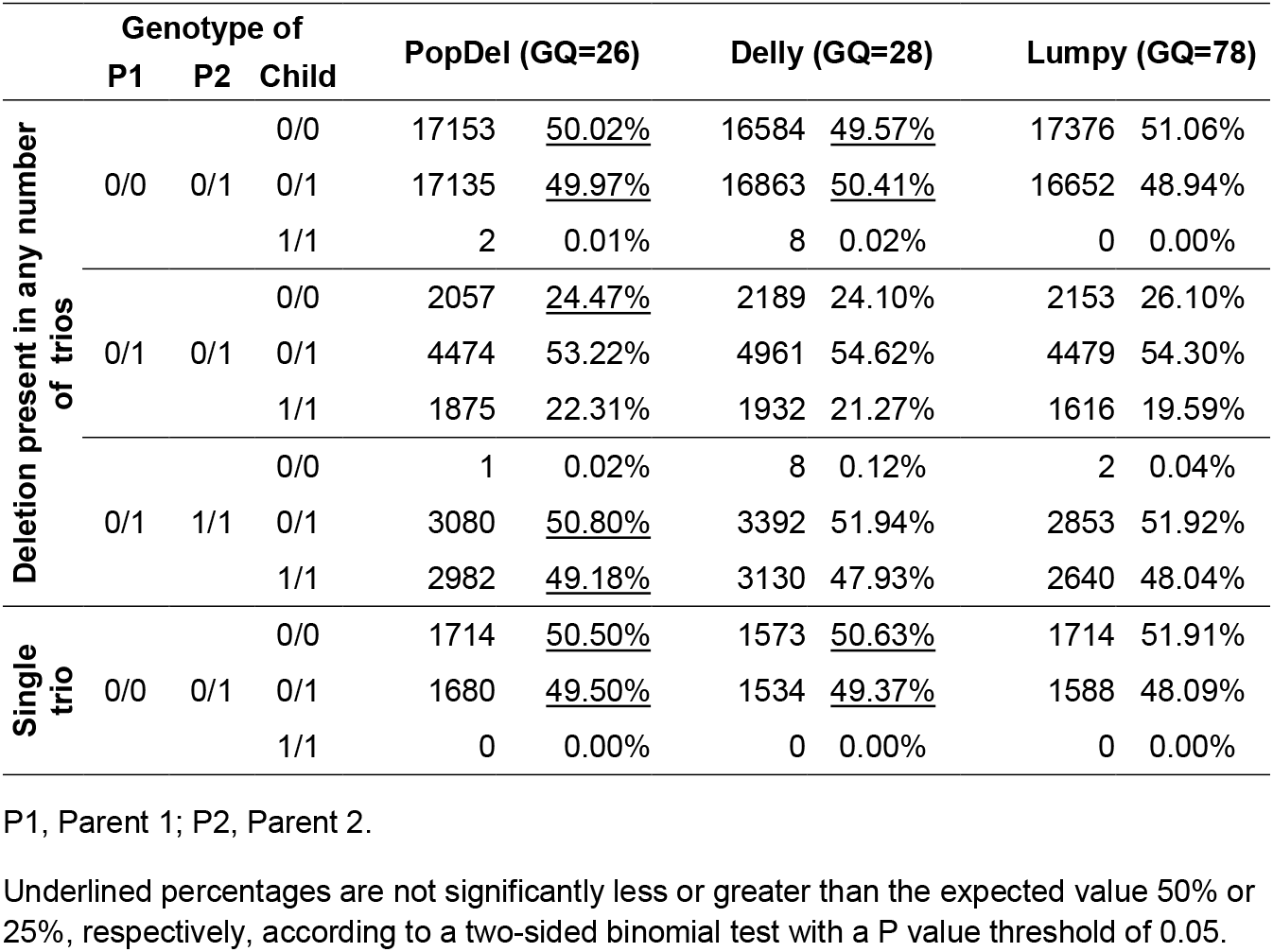
Transmission of deletions in the 49 Polaris trios after filtering for Hardy-Weinberg equilibrium (P value 0.01) and genotype quality (GQ) to a Mendelian inheritance error rate just below 0.3%.

**Table 2.** Running times (wall-clock time, CPU hours in parentheses) on NA12878 and the Polaris Diversity cohort.

|  | **NA12878**<br>**(single individual)** | **Polaris Diversity cohort**<br>**(150 individuals)** |
| --- | --- | --- |
| **PopDel** | 0:25 (0:58) | 56:17 (111:12) |
| **Delly** | 1:42 (1:40) | 389 :43 (371:27) * |
| **Lumpy (via Smoove)** | 0:18 (0:30) | 87:58 (103:21) * |
| **Manta** | 7:09 (7:09) | - |

\* Single sample calling with subsequent variant merging and sample-wise genotyping.
Note that Delly, Lumpy and Manta report other types of SVs apart from deletions. These SVs were excluded after single sample calling before merging to reduce running times.

### Application to population-scale data from Iceland

We applied PopDel to whole-genome data of 49,962 Icelanders including 6,794 parent-offspring trios (Methods, **Supplementary Text**). The average number of deletions PopDel reports per Icelander on the autosomes is 1504 heterozygous and 209 homozygous deletions (genotype quality threshold 25). The Mendelian inheritance error rate in the 6,794 trios is 1.4% (1,963 consistent deletions on average per trio). The transmission rate for 4,256 deletions unique to a single trio is 49.2%, which is again not significantly different from the expected 50% (two-sided binomial test, P value 0.32). This implies that the majority of errors appear as common deletions shared by several individuals.

### Identification of a *de novo* deletion in the Polaris data

We searched for *de novo* deletions (Methods) in the Polaris Kids cohort and identified twelve candidate *de novo* variants. Manual inspection suggests that three of them are true *de novo* events: a 8901 bp deletion at chr6:93035858-93044759 in the Spanish female HG01763 (Figure 6), an exonic 984 bp deletion at chr6:27132732-27133716 in the Spanish male HG01683 in the *H2BC11* gene, and an exonic 769 bp deletion at chr7:105505500-105506269 in the Chinese female HG00615 in the *PUS7* gene. The 8901 bp deletion in HG01763 is flanked and overlapped by SNVs that allow us to phase and confirm the *de novo* event. A SNV that overlaps with read pairs supporting the deletion indicate that the deletion haplotype was inherited from the mother (HG01762). Further evidence for this to be a true *de novo* event is given by 25 SNVs within the deletion that confirm the child to carry a single haplotype where both parents are heterozygous. All three individuals are heterozygous for numerous SNVs upstream of the deletion. Four of the SNVs within the deletion confirm that the event happened on a maternal haplotype. Given that this deletion is intergenic and HG01763 is part of a cohort of healthy individuals^1^, we expect the *de novo* deletion not to be of medical relevance. The closest transcript annotations in Gencode v29^40^ are the lncRNA *AL138731.1* at a distance of 25.6 kb and the *EPHA7* gene in a distance of 195.3 kb.

**Figure 6.**
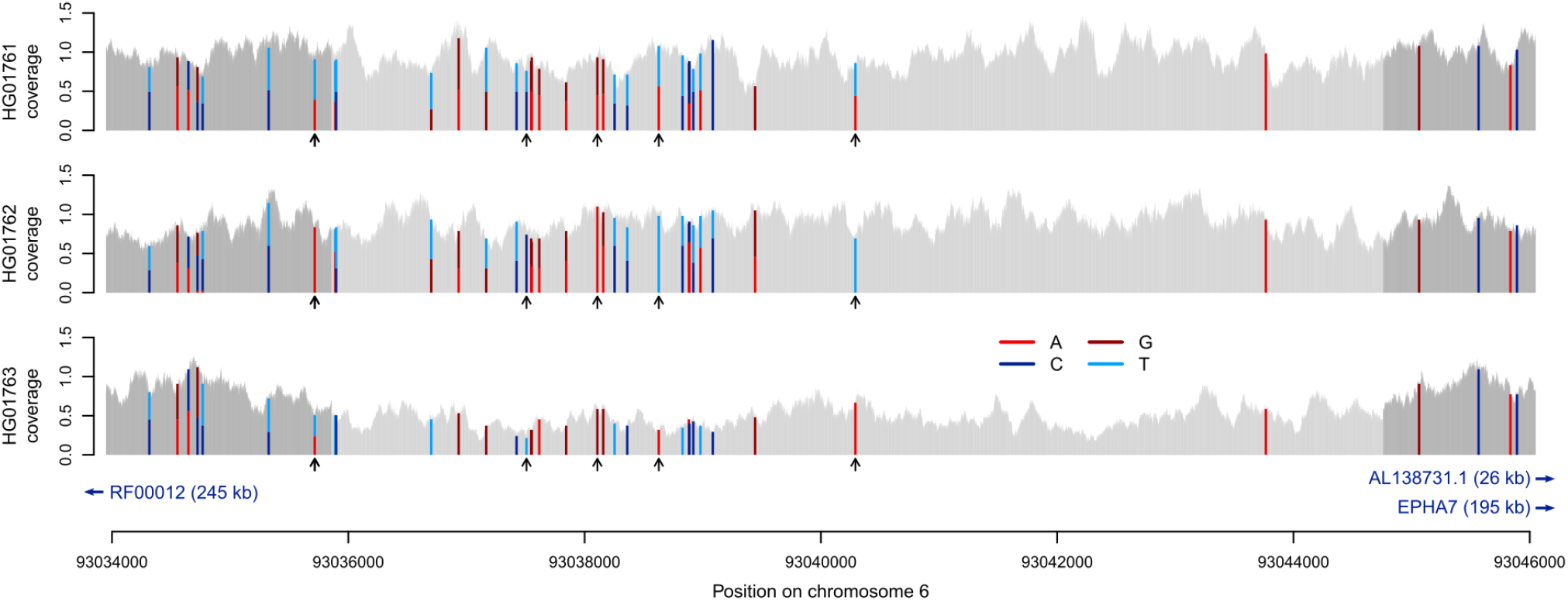
*De novo* deletion identified by PopDel in one individual of the Polaris Kids cohort. The child (HG01763, bottom) is carrier of a heterozygous deletion (indicated by the light gray interval) that is not present in either parent. Arrows mark SNPs that allow phasing of the haplotypes in the child.

### Running times on public benchmarking data

We assessed the running time of PopDel compared to Delly and Lumpy via Smoove on the data of the NA12878 genome and the 150 genomes in the Polaris Diversity cohort (Methods, Table 1). With a total wall clock running time of approximately 25 minutes for profile creation and deletion calling of the NA12878 genome, PopDel is almost as fast as Lumpy via the Smoove pipeline and 4 times faster than Delly. A similar behavior can be observed on the 150 genomes of the Polaris Diversity cohort confirming scalability: PopDel completes deletion calling within less than 2 days and 9 hours, which is similar to the running time of Lumpy via Smoove (3 days and 16 hours) and several times faster than Delly (16 days, 6 hours). PopDel can be trivially parallelized by creating profiles of different individuals in parallel and splitting the joint calling by genomic region.

## Discussion

Identification and genotyping of structural variation in large sequencing cohorts is a major computational challenge. To enable the joint analysis of the increasingly large cohorts that are being sequenced, we developed a novel deletion calling approach implemented in the tool PopDel. Compared to existing approaches, the joint calling approach in PopDel greatly simplifies the analysis workflow, shows comparable if not better accuracy and has a competitive running time. PopDel scales to very large cohorts as our tests on population-scale data from Iceland substantiates.

PopDel has high accuracy independent of the number of jointly analyzed individuals and across the deletion allele frequency spectrum. On data of a single genome, PopDel shows higher recall than other tools at high precision. On the Polaris Diversity cohort, the deletions called by PopDel recapitulate previous population genetic results showing that Africans carry on average more deletions than other continental groups and confirming that joint calling can be used to identify population structure. On the Polaris Kids cohort, PopDel identifies more deletions at a better transmission rate compared to other tools and reports a *de novo* deletion of about 9 kb. On Icelandic data, PopDel identifies deletions jointly in almost 50,000 genomes maintaining an excellent transmission rate for rare variants. All results confirm that the joint calling approach in PopDel is accurate across the allele frequency spectrum and the number of individuals analyzed.

The *de novo* deletion in the Polaris Kids cohort together with the good transmission rate of rare variants in the large number of Icelandic genomes demonstrates that PopDel’s joint calling approach provides a basis for studying rarely observed *de novo* deletion events. A previous study^41^ verified in 258 healthy trios seven *de novo* deletions that fall into the size range addressed by us. Given their rate of *de novo* deletions, we expect to observe 1.33 *de novo* deletions of medium size in the 49 Polaris trios. This is well in line with our finding of three candidate *de novo* events including one that we could confirm based on nearby SNVs.

When we tested an early version of PopDel on a selected 54 kb region covering the *LDLR* gene in 43,202 Icelanders, we identified a previously unknown 2.5 kb deletion in three closely related Icelanders shown to affect LDL levels (Björnsson et al. manuscript in preparation). This finding shows that PopDel is able to identify variants of biomedical interest even if they are present at a very low allele frequency in a population-scale cohort, and showcases the importance of SVs in human health.

PopDel consists of two steps: creation of read pair profiles per individual and joint deletion calling. The computational advantage of this two-step design is that the large input BAM files containing aligned read data need to be processed only once. The joint calling step takes the information needed for deletion detection and genotyping from the small read pair profiles. This implies that additional genomes, “the N+1^st^ genome”, can be added to the analysis without the need to access all input BAM files reducing the computational burden considerably. PopDel is currently limited to deletions. However, the two-step design and the likelihood ratio test can be generalized to junctions of other types of SVs. We are aiming for extending our approach accordingly in the near future.

## Code and data availability

PopDel is available at https://github.com/kehrlab/PopDel (v1.2.2, GNU GPLv3 license). The generated VCF files used for evaluation of all simulated and real data sets are available at Zenodo (DOI: 10.5281/zenodo.3992607).

## URLs

Smoove pipeline, https://github.com/brentp/smoove; Long read reference call set for HG001, ftp://ftp-trace.ncbi.nlm.nih.gov/giab/ftp/data/NA12878/NA12878_PacBio_MtSinai/NA12878.sorted.vcf.gz; Short read reference call set for HG001, ftp://ftp-trace.ncbi.nlm.nih.gov/giab/ftp/technical/svclassify_Manuscript/Supplementary_Information/Personalis_1000_Genomes_deduplicated_deletions.bed; Preliminary variant set for HG002, ftp://ftp-trace.ncbi.nlm.nih.gov/giab/ftp/data/AshkenazimTrio/analysis/NIST_SVs_Integration_v0.6/HG002_SVs_Tier1_v0.6.vcf.gz; High-confidence regions for HG002, ftp://ftp-trace.ncbi.nlm.nih.gov/giab/ftp/data/AshkenazimTrio/analysis/NIST_SVs_Integration_v0.6/HG002_SVs_Tier1_v0.6.bed; NCBI genome remapping service, https://www.ncbi.nlm.nih.gov/genome/tools/remap;

## Author contributions

B.K. and B.V.H. conceived the approach. S.N. and B.K. developed PopDel, designed the experiments and evaluated the results. S.N. simulated data, performed analyses on simulated and public data and tested on Icelandic data. B.V.H. applied PopDel on Icelandic data and assisted in testing. E.B., P.S. and K.S. studied the *LDLR* deletion. H.J assisted in the computation of transmission rates. H.P.E. and D.B. assisted in the analyses of Icelandic data. B.K and S.N. drafted the manuscript with feedback from B.V.H. All authors revised the draft and approved the final manuscript.

## Methods

### Read pair profile creation

PopDel reduces a coordinate-sorted BAM file of each sample into a read pair profile in a custom binary format (**Supplementary Text**). This profile stores positions and insert sizes of read pairs that align confidently (**Supplementary Table 1**) to the reference genome. In addition, the profile file contains meta information, including a distribution of insert sizes across the sample and an index, which allows for jumping to genomic positions in the profile. We define the insert size as the distance between the leftmost alignment position of the forward read to the rightmost alignment position of the reverse read in the pair extended by any clipped bases (**Supplementary Figure 1**). The null distribution of insert sizes is estimated by sampling the BAM file using pre-defined but user-configurable genomic regions with good mappability (**Supplementary Text**). If more than one library has been sequenced for a sample, PopDel writes separate profile data per read group to the profile file. An excerpt of an example profile is shown in **Supplementary Table 2**. The profiling vastly reduces I/O during joint calling as the size of the profiles is on average only 1.76% of the original BAM file size (**Supplementary Figure 2**).

### Likelihood ratio test for joint deletion calling

For a given genomic window, our likelihood ratio test compares the relative likelihood that a deletion of a particular length *l* overlaps the window, against the relative likelihood of observing the reference haplotype:

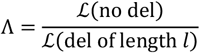

Our null hypothesis is that the data observed in a window is drawn from the reference model (numerator) rather than the deletion model (denominator). We reject the null hypothesis in PopDel using a cutoff for Λ calculated from −2 logΛ ~ *χ*^2^ with a p-value threshold of 0.01 (one-tailed) and 1 degree of freedom in order to decide if the window overlaps a deletion of length *l*.

Let 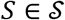 be a single sample from the set of all samples 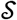 and let *I^s^* be the list of insert sizes for all the read pairs of *S* overlapping the given window (**Supplementary Figure 1**). Furthermore, let Δ^S^= (*i-−μ_S_* |*i ∈ I^S^*) be the deviations of the insert sizes from the mean insert size of the sample *μ_S_*. We assume independence of samples and calculate the relative likelihood of the reference model as the product of the samples’ likelihoods 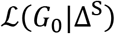 for the reference genotype *G*_0_

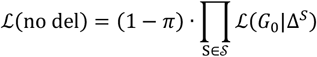

where *π* is the prior probability of observing a deletion (default 10^−4^). For the likelihood of the deletion model we use the weighted sums of all three genotype likelihoods in a similar product

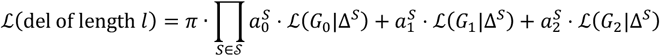

where the 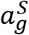 are sample- and genotype-specific weights (see below) with genotypes *g* ∈ {0,1,2} corresponding to 0, 1 or 2 variant alleles and 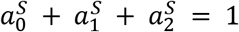 for any 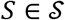.

### Iterative estimation of parameters for the likelihood ratio test

The likelihood ratio test requires as input a deletion length *l*, genotype likelihoods 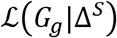 for all samples *S* and sample- and genotype-specific weights 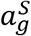. PopDel estimates these values for each window iteratively from the profiles together with an allele frequency *f* that is needed for updating the weights (Figure 1). For simplicity, the following assumes one read group per sample. Our implementation in PopDel also handles multiple read groups (**Supplementary Text**). To be able to detect deletions of different lengths from different haplotypes overlapping the same window, the iteration and likelihood ratio test are performed for several initializations of the deletion length. Initial lengths are estimated by identifying samples with similar third quartiles of Δ^*S*^ via greedy clustering (**Supplementary Text**). The initial allele frequencies *f* are set to the fraction of deletion-supporting read pairs of all samples in the window (**Supplementary Text**). To calculate the genotype likelihoods of the three genotypes *G*_0_, *G*, and *G*_2_ of a single sample *S*, PopDel transforms the insert size histogram of *S* to reflect how many read pairs with a given insert size deviation *δ* ∈ Δ^*S*^ are expected to overlap a window of size *w* (**Supplementary Text**). We denote the resulting relative likelihood of observing a read pair with insert size deviation *δ* as *H^S^*(*δ*).

PopDel calculates the likelihoods 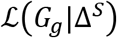 as

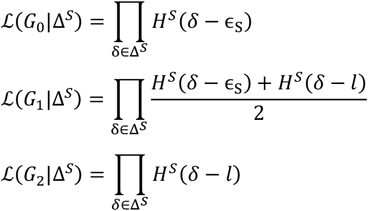

where *ϵ_S_* is a sample-specific reference shift (**Supplementary Text**) that accounts for local biases of the data such as GC-content^42,43^

Our *sample- and genotype-specific weights* 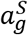 are designed to give low weight to samples with a small likelihood for the genotype and a high weight to those with a good likelihood for the genotype. Furthermore, the weights make it more likely to observe a carrier genotype when the allele frequency is high:

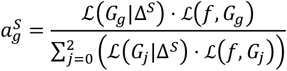

with 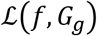 as the expected genotype frequencies given the population allele frequency *f*:

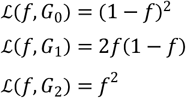

Given the weights, we update the *allele frequency f* using:

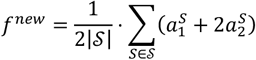

To update the *deletion length l*, probabilities 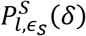 reflecting that a given insert size deviation *δ* resulted from a distribution shifted by *l* rather than by *ϵ_S_* are calculated as

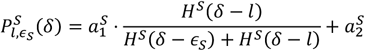

and used to update *l* jointly across all samples as the weighted sum over all insert size deviations:

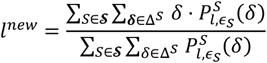

The iteration for parameter estimation terminates when both the allele frequency and deletion length converge or additional termination conditions are met (**Supplementary Text**), e.g. reaching the maximum number of iterations (default 15). A *start position* of the potential deletion is estimated during above calculations by keeping track of the rightmost aligned positions of the forward reads of read pairs whose *δ* support the deletion estimate (**Supplementary Text**).

### Combining consecutive deletion windows

The likelihood ratio tests are performed per initialized deletion length for each genomic window (of size 30 bp by default). The deletions identified by PopDel typically overlap several consecutive windows and in each window the null hypothesis of the likelihood ratio test may be rejected. To provide the user with a non-redundant list of deletion variants, adjacent windows that support the same deletion are combined. PopDel sorts all pairs of windows and deletion lengths, for which the null hypothesis of the likelihood ratio test can be rejected, in ascending order of the predicted deletion start position, deletion length and deletion likelihood ratio. Traversing this sorted list of windows *w*_0_, *w*_1_,…, a window *w_i_, i* ≥ 0 is combined with another window *w_i+k_, k* > 0 if their start positions and deletion sizes are similar enough (**Supplementary Text**). When no more windows can be combined with *w_i_*, a deletion is output with a start position and length calculated as the median over all combined windows. The algorithm continues with the next window *w*_*i+k*+1_ that has not been combined with any other window so far.

### Deletion output

We report the mean PHRED-scaled genotype likelihoods across the combined windows of one sample in the output. Samples without sufficient data or much higher than average coverage at the locus are not genotyped (**Supplementary Text**). The allele frequency is estimated by counting the number of alleles predicted to carry the variant, divided by the total number of genotyped alleles. We report a genotype quality as the difference of the best and second best PHRED-scaled genotype likelihoods.

### Simulation of sequencing data with deletions

We simulated two cohorts of sequencing data, the first consisting of 1,000 diploid individuals carrying random deletions and the second consisting of 500 individuals carrying deletions reported in the 1000 Genomes Project (G1k). For the first cohort, we simulated a set of 2,000 deletion variants with uniformly distributed lengths between 100 and 10,000 bp, uniformly distributed allele frequencies between 0 and 1, and uniformly distributed positions on chromosome 21 of GRCh38. Regions containing ‘N’s were excluded and deletion were required to be at least 1000 bp apart. For the second cohort, we downloaded deletions identified in the 1000 Genomes Project (see URLs) on chromosomes 17 to 22 of GRCh37 and their reported allele frequencies.

Using the random deletion set, we created 2,000 haplotypes by sampling deletions according to their allele frequency and inserting them into GRCh38. Using the G1k deletions, we created 1,000 haplotypes by sampling deletions according to their allele frequencies and inserting them into GRCh37. The haplotypes were combined into 1,000 and 500 diploid samples, respectively. These samples were used to simulate NGS reads with *art_illumina*^44^ and the reads aligned to GRCh38 or GRCh37 using *BWA-mem*^45^.

### Setup of SV callers on simulated data

PopDel (1.2.2) and Manta (1.6.0) were run with an option to limit the calling to chromosome 21. Smoove (0.2.4 with Lumpy 0.2.13 and SVtyper 0.7.0) was applied as recommended by its authors using the provided exclude regions for GRCh38 and GRCh37, and excluding mappings not on the simulated chromosomes. Delly (0.7.8) was applied without small indel realignment (option -n). GRIDSS (1.8.1) was provided a maximum heap size of 8 GB. All tools were applied on increasing numbers of BAM files (up to 1,000 or until failure) in steps of 1 up to 10, steps of 10 up to 100, and steps of 100 up to 1,000.

### Evaluation on simulated data

Running time and memory consumption were measured on a dedicated workstation (Intel Xeon E5-1630v3 8×3.5GHz, 64GB RAM) using the Unix time command. For tools consisting of multiple steps, the running times are the sum of the time taken by all steps from input BAM files to output VCF file. The memory consumption is stated as the maximum memory consumption of all steps. As GRIDSS produces two break-ends per deletion, corresponding pairs of break-ends were collapsed into a single call and “LOW_QUAL” variants were removed. The calls of Delly were not filtered for variants that have the filter field set to “PASS” as this has a negative impact on its performance. A call is considered to match a simulated variant in case of a reciprocal overlap of at least 50%. Each simulated variant is allowed to be matched with only one predicted variant. See **Supplementary Text** for results using alternative match criteria.

### Accession numbers and reference call sets for HG001 and HG002

We used short read data for the HG001/NA12878 genome available under ERA run accession ERR194147, and a subset of the short read data for the HG002/NA24385 genome and parental genomes HG003/NA24149 and HG004/NA24143 available under BioProject accessions PRJNA200694. The precise list of run accession numbers for HG002 and his parents is given in the **Supplementary Text**.

GiaB short read and PacBio reference call sets were downloaded (see URLs), filtered for deletion variants. Liftover from GRCh37 to GRCh38 was performed for HG001/NA12878 using the *NCBI Genome Remapping Service* (see URLs). All variants on contigs apart from chromosomes 1 to 22 and chromosome X were removed and VCF-to-BED conversion was performed for the PacBio call set.

### Sample preparation and setup of SV callers on GiaB and Polaris data

All samples were aligned to the human reference genome GRCh38^46^ using *BWA-mem*^47,48^ except for the data of HG002/NA24385 and his parents, which was aligned to GRCh37. All SV callers were applied as recommended by the authors and, if possible, limited to the reference sequence of the 22 autosomes. Delly was run with the option -n. All tools could be run jointly on the GiaB trio data. Smoove, the recommended pipeline for running Lumpy, recommends a workflow consisting of single-sample calling, merging and sample-wise re-genotyping for cohorts of 40 or more individuals and was applied accordingly. Joint calling with Delly and Manta did not finish within 4 weeks on the Polaris data. Therefore, Delly was run using single-sample calling with merging and sample-wise re-genotyping following the germline SV calling workflow described by the authors (**Supplementary Text**). For Manta, we could not find a description of a similar workflow, so we excluded it from the analysis on the Polaris data. Calls other than deletions were removed as early as possible in Delly’s and Smoove’s workflows to reduce running time (**Supplementary Text**). The sample order of the Polaris Diversity and Kids cohorts was shuffled but the same for all tools and no tool was provided pedigree information.

### Variant filtering on real data

Deletions found in HG002/NA24385 and his parents were filtered using the high-confidence regions prepared by the GiaB consortium (see URLs). All other deletion sets were filtered for centromeric regions. Centromeric regions were obtained through the UCSC table browser (“group: Mapping and Sequencing”, “track: Centromeres”) for GRCh38. Any deletions in a reference set or call set having any overlap with a centromeric region or a region outside of the high-confidence regions were removed. Overlap was determined using *bedtools intersect*^49^ (**Supplementary Text**).

Only deletions on the 22 autosomes were considered for analysis. Deletions were filtered to the size range from 500 to 10,000 bp. Two deletions were considered the same if they had a reciprocal overlap of 50% or more. The Polaris data was filtered to high-confidence deletions with genotype quality scores above a fixed threshold. This threshold was chosen once per tool on the Polaris Kids cohort such that the Mendelian inheritance error rate dropped below 0.3%^38^: 26 for PopDel, 28 for Delly and 78 for Lumpy. To search for *de novo* deletions, the genotype threshold for PopDel was set to 50 and all twelve resulting candidate deletions were inspected manually in IGV^50^. In all real data analyses, Delly variants were only considered if they had the ‘FILTER’ field set to ‘PASS’.

### Principal Component Analysis

Predicted genotypes of the Polaris Diversity cohort were converted into a variant/sample matrix containing deletion allele counts. Uninformative deletions and those in linkage disequilibrium were removed (**Supplementary Text**). PCA was computed using the R-function *prcomp*.

### Mendelian inheritance error rate and transmission rate

All deletions that are not in Hardy-Weinberg equilibrium (P value threshold 0.01) were removed. For all reported deletions, the three genotypes in each trio were inspected for Mendelian consistency (**Supplementary Table 3**). Trios with one or more missing genotypes and trios with all three samples genotyped as 0/0 were ignored. The transmission rate was calculated as the number of deletion alleles transmitted from the heterozygous parents to the children divided by the number of considered deletions. If indicated, we calculated the transmission rate considering only deletions that were called in a single trio, where one parent is a heterozygous carrier and the other parent carries the reference allele on both haplotypes.

### Sequence data from 49,962 Icelanders

DNA was isolated from both blood and buccal samples. All participating subjects signed informed consent. The personal identities of the participants and biological samples were encrypted by a third-party system approved and monitored by the Data Protection Authority. The National Bioethics Committee and the Data Protection Authority in Iceland approved these studies. The Icelandic samples were whole-genome sequenced at deCODE Genetics using Illumina GAIIx, HiSeq, HiSeqX and NovaSeq sequencing machines, and sequences were aligned to the human reference genome (GRCh38)^46^ using *BWA-mem*^47,48^. Details of the sample preparation, paired-end sequencing, read processing and alignment, and selection of the final set of BAM files have been previously described^51^.

### Running time measurements on real data

The running time on NA12878 was measured using the Unix time command on a dedicated workstation (Intel Xeon E5-1630v3 8×3.5GHz, 64GB RAM). Due to limited storage capacities on our workstations, running times for the Polaris Diversity cohort were measured on the BIH high-performance compute cluster as reported in SGE log files.

