## Supplementary Material for "PopDel identifies medium-size deletions jointly in tens of thousands of genomes"

### PopDel calls medium-sized deletions jointly in tens of thousands of genomes

#### Supplementary tables

#### Supplementary tables

Supplementary table 1 - Alignment requirements for read pairs to be included in PopDel's profiles.

| Parameter | Description | Default* |
|---|---|---|
| orientation | The two reads of the pair align in forward-reverse orientation. | true |
| unclipped | Minimum number of aligned base pairs per read. | 50 |
| mapping-qual | Minimum value of the mapping quality per read. | 1 |
| align-score | Minimum alignment score per read (in percent of the length of the aligned part of the read). | 80 |
| flags-unset | SAM flags that are not set in both reads | 3840 |

* All parameter values are user configurable.

Supplementary table 2 - Human readable representation of PopDel's profile format.

| Chromosome | Window position | Read pair information |
|---|---|---|
| chr21 | 5035777 | 230:-10, 233:-34 |
| chr21 | 5036033 | 3:-37, 11:0, 19:-3, 214:15 |
| chr21 | 5036289 | 4:-85, 4:-29, 13:-96, 13:-78:17, 80:-5 |
| chr21 | 5036545 | 172:5, 178:51, 180:13, 192:19, 248:-58 |
| chr21 | 5036801 | 51:35, 62:-50, 83:-60, 85:33, 110:88, 218:104 |
| chr21 | 5037057 | 196:53, 210:12, 210:43, 232:-77, 253:-19 |
| chr21 | 5037313 | 11:51, 15:-63, 27:77, 44:10, 64:-73, 65:-88 |

The format is divided into windows of 256 bp. The start position of each window is given in column one and two. The information on read pairs that fall in the respective window is given in column three, divided by ','. The number before the ':' indicates the offset of the start position from the start of the window and the value after the ':' is the deviation from the median insert size μ. The start position of a pair refers to the rightmost aligned base of the forward read of a pair. Each read pair is only listed in the window that contains its start position.

Supplementary table 3 - Visualization of Mendelian consistency.

| | Parent A: 0/0 | | | Parent A: 0/1 | | | Parent A: 1/1 | | |
|---|---|---|---|---|---|---|---|---|---|
| Parent B: 0/0 | 0/0 | 0/1 | 1/1 | 0/0 | 0/1 | 1/1 | 0/0 | 0/1 | 1/1 |
| Parent B: 0/1 | | | | 0/0 | 0/1 | 1/1 | 0/0 | 0/1 | 1/1 |
| Parent B: 1/1 | | | | | | | 0/0 | 0/1 | 1/1 |

Mendelian inheritance error

Consistent with Mendelian inheritance, uninformative for transmission rate

Consistent with Mendelian inheritance, one possible transmission

Consistent with Mendelian inheritance, two possible transmissions

The outer fields indicate the genotypes of the parents while the numbers of the colored fields indicate the observable genotypes of the child. The colors indicate if the genotype of the child Is consistent with the rules of Mendelian inheritance or not. For the calculation of the transmission rate only the informative fields are considered.

Supplementary table 4 – Number of true positives calls on deletions simulated randomly on human chromosome 21.

| Samples | PopDel | Delly | Manta | Lumpy | GRIDSS |
|---:|---:|---:|---:|---:|---:|
| 1 | 1157 | **1166** | 1145 | 1140 | 1148 |
| 2 | **1414** | 1408 | 1395 | 1380 | 1396 |
| 3 | **1530** | 1517 | 1509 | 1491 | 1515 |
| 4 | **1587** | 1583 | 1572 | 1553 | 1579 |
| 5 | **1633** | 1630 | 1616 | 1596 | 1625 |
| 6 | **1652** | 1645 | 1633 | 1611 | 1643 |
| 7 | **1672** | 1664 | 1652 | 1629 | 1663 |
| 8 | **1690** | 1678 | 1666 | 1643 | 1679 |
| 9 | **1696** | 1683 | 1671 | 1648 | 1685 |
| 10 | **1702** | 1687 | 1676 | 1653 | 1690 |
| 20 | **1748** | 1732 | 1723 | 1702 | 1736 |
| 30 | **1766** | 1748 | 1739 | 1718 | 1749 |
| 40 | **1775** | 1757 | 1746 | 1728 | 1756 |
| 50 | **1781** | 1763 | 1754 | 1734 | 1764 |
| 60 | **1783** | 1765 | 1755 | 1737 | 1767 |
| 70 | **1787** | 1767 | 1758 | 1739 | 1771 |
| 80 | **1789** | 1769 | 1759 | 1740 | 1771 |
| 90 | **1790** | 1770 | 1761 | 1741 | 1770 |
| 100 | **1790** | 1771 | 1762 | 1742 | 1772 |
| 200 | **1794** | 1777 | 1761 | 1748 | 1773 |
| 300 | **1794** | 1782 | 1761 | 1750 | 1777 |
| 400 | **1795** | 1785 | 1764 | 1751 | 1779 |
| 500 | **1795** | 1787 | 1763 | 1751 | NA |
| 600 | **1793** | 1790 | 1764 | 1753 | NA |
| 700 | **1795** | 1792 | 1765 | 1755 | NA |
| 800 | **1795** | 1793 | 1765 | 1755 | NA |
| 900 | **1795** | 1794 | 1762 | 1755 | NA |
| 1000 | **1794** | 1793 | 1762 | 1756 | NA |

The maximum value of each row is highlighted.

Supplementary table 5 - Number of false positive calls on deletions simulated randomly on human chromosome 21.

| Samples | PopDel | Delly | Manta | Lumpy | GRIDSS |
|---|---|---|---|---|---|
| 1 | **0** | **0** | **0** | **0** | 0 |
| 2 | **0** | **0** | **0** | **0** | 0 |
| 3 | **0** | **0** | **0** | **0** | 0 |
| 4 | **0** | **0** | **0** | **0** | 0 |
| 5 | **0** | **0** | **0** | **0** | 0 |
| 6 | **0** | **0** | **0** | **0** | 0 |
| 7 | **0** | **0** | **0** | **0** | 0 |
| 8 | **0** | **0** | 1 | **0** | 0 |
| 9 | **0** | **0** | 1 | **0** | 0 |
| 10 | **0** | **0** | 1 | **0** | 0 |
| 20 | **0** | **0** | **0** | **0** | 0 |
| 30 | **0** | **0** | **0** | **0** | 0 |
| 40 | 1 | **0** | 1 | **0** | 0 |
| 50 | 1 | **0** | 1 | **0** | 5 |
| 60 | **0** | **0** | 1 | **0** | 9 |
| 70 | **0** | **0** | 1 | **0** | 17 |
| 80 | **0** | **0** | 1 | **0** | 29 |
| 90 | **0** | 1 | 1 | **0** | 58 |
| 100 | 1 | 1 | 1 | **0** | 69 |
| 200 | **0** | 2 | 4 | **0** | 82 |
| 300 | 1 | 3 | 4 | **0** | 134 |
| 400 | **0** | 5 | 4 | **0** | 309 |
| 500 | 1 | 5 | 3 | **0** | NA |
| 600 | 2 | 6 | 4 | **0** | NA |
| 700 | 1 | 8 | 4 | **0** | NA |
| 800 | 1 | 11 | 4 | **0** | NA |
| 900 | 1 | 13 | 4 | **0** | NA |
| 1000 | 1 | 13 | 4 | **0** | NA |

The minimum values of each row are highlighted.

Supplementary table 6 - Number of false negatives on deletions simulated randomly on human chromosome 21.

| Samples | PopDel | Delly | Manta | Lumpy | GRIDSS |
|---|---|---|---|---|---|
| 1 | 159 | **150** | 171 | 176 | 168 |
| 2 | **185** | 191 | 204 | 219 | 203 |
| 3 | **189** | 202 | 210 | 228 | 204 |
| 4 | **204** | 208 | 219 | 238 | 212 |
| 5 | **205** | 208 | 222 | 242 | 213 |
| 6 | **203** | 210 | 222 | 244 | 212 |
| 7 | **202** | 210 | 222 | 245 | 211 |
| 8 | **201** | 213 | 225 | 248 | 212 |
| 9 | **200** | 213 | 225 | 248 | 211 |
| 10 | **198** | 213 | 224 | 247 | 210 |
| 20 | **203** | 219 | 228 | 249 | 215 |
| 30 | **197** | 215 | 224 | 245 | 214 |
| 40 | **197** | 215 | 226 | 244 | 216 |
| 50 | **198** | 216 | 225 | 245 | 215 |
| 60 | **199** | 217 | 227 | 245 | 215 |
| 70 | **197** | 217 | 226 | 245 | 213 |
| 80 | **198** | 218 | 228 | 247 | 216 |
| 90 | **198** | 218 | 227 | 247 | 218 |
| 100 | **199** | 218 | 227 | 247 | 217 |
| 200 | **199** | 216 | 232 | 245 | 220 |
| 300 | **202** | 214 | 235 | 246 | 219 |
| 400 | **203** | 213 | 234 | 247 | 219 |
| 500 | **203** | 211 | 235 | 247 | NA |
| 600 | **205** | 208 | 234 | 245 | NA |
| 700 | **204** | 207 | 234 | 244 | NA |
| 800 | **204** | 206 | 234 | 244 | NA |
| 900 | **204** | 205 | 237 | 244 | NA |
| 1000 | **206** | 207 | 238 | 244 | NA |

The minimum values of each row are highlighted.

Supplementary table 7 – Number of true positive calls for 1000 Genomes Project deletions introduced into simulated chromosomes 17 to 22.

| Samples | PopDel | Delly | Manta | Lumpy |
|---|---|---|---|---|
| 1 | 176 | 163 | 178 | **192** |
| 2 | 290 | 262 | 287 | **303** |
| 3 | 379 | 343 | 369 | **388** |
| 4 | 417 | 378 | 410 | **427** |
| 5 | 456 | 408 | 456 | **470** |
| 6 | 491 | 441 | 492 | **505** |
| 7 | 520 | 466 | 528 | **537** |
| 8 | 541 | 483 | 551 | **559** |
| 9 | 558 | 502 | 569 | **581** |
| 10 | 582 | 524 | 592 | **604** |
| 20 | 773 | 717 | 790 | **803** |
| 30 | 913 | 855 | 938 | **952** |
| 40 | 999 | 948 | 1025 | **1042** |
| 50 | 1085 | 1040 | 1108 | **1127** |
| 60 | 1159 | 1126 | 1193 | **1215** |
| 70 | 1272 | 1248 | 1304 | **1331** |
| 80 | 1339 | 1313 | 1371 | **1400** |
| 90 | 1401 | 1371 | 1435 | **1464** |
| 100 | 1466 | 1443 | 1502 | **1534** |
| 200 | 2048 | 2029 | 2088 | **2138** |
| 300 | 2452 | 2431 | 2442 | **2559** |
| 400 | 2721 | 2699 | 2639 | **2846** |
| 500 | 2962 | 2944 | 2856 | **3111** |

The maximum value of each row is highlighted.

Supplementary table 8 - Number of false positive calls for 1000 Genomes Project deletions introduced into simulated chromosomes 17 to 22.

| Samples | PopDel | Delly | Manta | Lumpy |
|---:|---:|---:|---:|---:|
| 1 | **0** | **0** | **0** | **0** |
| 2 | **0** | 1 | **0** | **0** |
| 3 | **0** | 1 | 1 | **0** |
| 4 | **0** | 1 | 1 | **0** |
| 5 | **0** | 1 | 1 | **0** |
| 6 | **0** | 1 | 1 | **0** |
| 7 | **0** | 1 | 1 | 1 |
| 8 | **0** | 1 | 1 | 1 |
| 9 | **1** | **1** | **1** | **1** |
| 10 | 2 | **1** | **1** | **1** |
| 20 | 2 | 1 | 7 | **0** |
| 30 | 5 | 1 | 8 | **0** |
| 40 | 4 | 2 | 9 | **0** |
| 50 | 4 | 1 | 8 | **0** |
| 60 | 5 | 3 | 12 | **0** |
| 70 | 9 | 3 | 13 | **0** |
| 80 | 8 | 3 | 15 | **0** |
| 90 | 8 | 4 | 16 | **0** |
| 100 | 8 | 4 | 16 | **0** |
| 200 | 13 | 10 | 18 | **0** |
| 300 | 16 | 13 | 20 | **0** |
| 400 | 13 | 17 | 24 | **0** |
| 500 | 15 | 26 | 28 | **0** |

The minimum values of each row are highlighted

Supplementary table 9 - Number of false negatives for 1000 Genomes Project deletions introduced into simulated chromosomes 17 to 22.

| Samples | PopDel | Delly | Manta | Lumpy |
|---|---|---|---|---|
| 1 | 35 | 48 | 33 | **19** |
| 2 | 42 | 70 | 45 | **29** |
| 3 | 46 | 82 | 56 | **37** |
| 4 | 48 | 87 | 55 | **38** |
| 5 | 47 | 95 | 47 | **33** |
| 6 | 47 | 97 | 46 | **33** |
| 7 | 51 | 105 | 43 | **34** |
| 8 | 54 | 112 | 44 | **36** |
| 9 | 59 | 115 | 48 | **36** |
| 10 | 60 | 118 | 50 | **38** |
| 20 | 86 | 142 | 69 | **56** |
| 30 | 100 | 158 | 75 | **61** |
| 40 | 110 | 161 | 84 | **67** |
| 50 | 112 | 157 | 89 | **70** |
| 60 | 129 | 162 | 95 | **73** |
| 70 | 138 | 162 | 106 | **79** |
| 80 | 140 | 166 | 108 | **79** |
| 90 | 144 | 174 | 110 | **81** |
| 100 | 150 | 173 | 114 | **82** |
| 200 | 202 | 221 | 162 | **112** |
| 300 | 239 | 260 | 249 | **132** |
| 400 | 255 | 277 | 337 | **130** |
| 500 | 284 | 302 | 390 | **135** |

The minimum value of each row is highlighted.

#### Supplementary figures

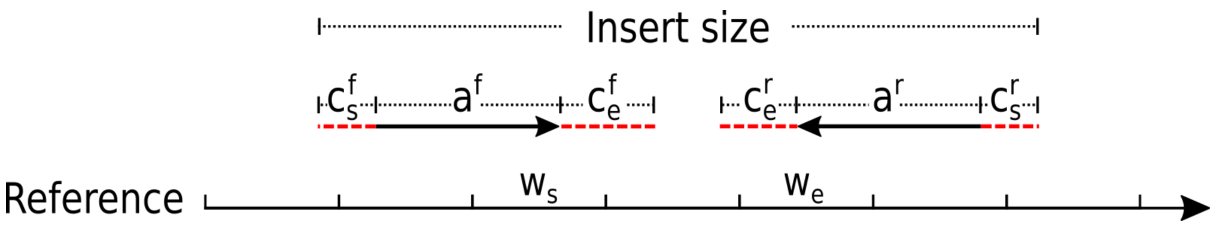

Supplementary figure 1 - Visualization of the parts of a read pair and its insert size. The insert size includes the length of the aligned parts of the reads ($a^f$, $a^r$) plus the outer soft clipped bases ($c_s^f$, $c_s^r$). The starting window $w_s$ is the window on the reference where the right end of $a^f$ aligns. The end window $w_e$ is the window where the left end of $a^r$ aligns. All windows between $w_s$ and $w_e$ (inclusive) are considered to be overlapped by the read pair.

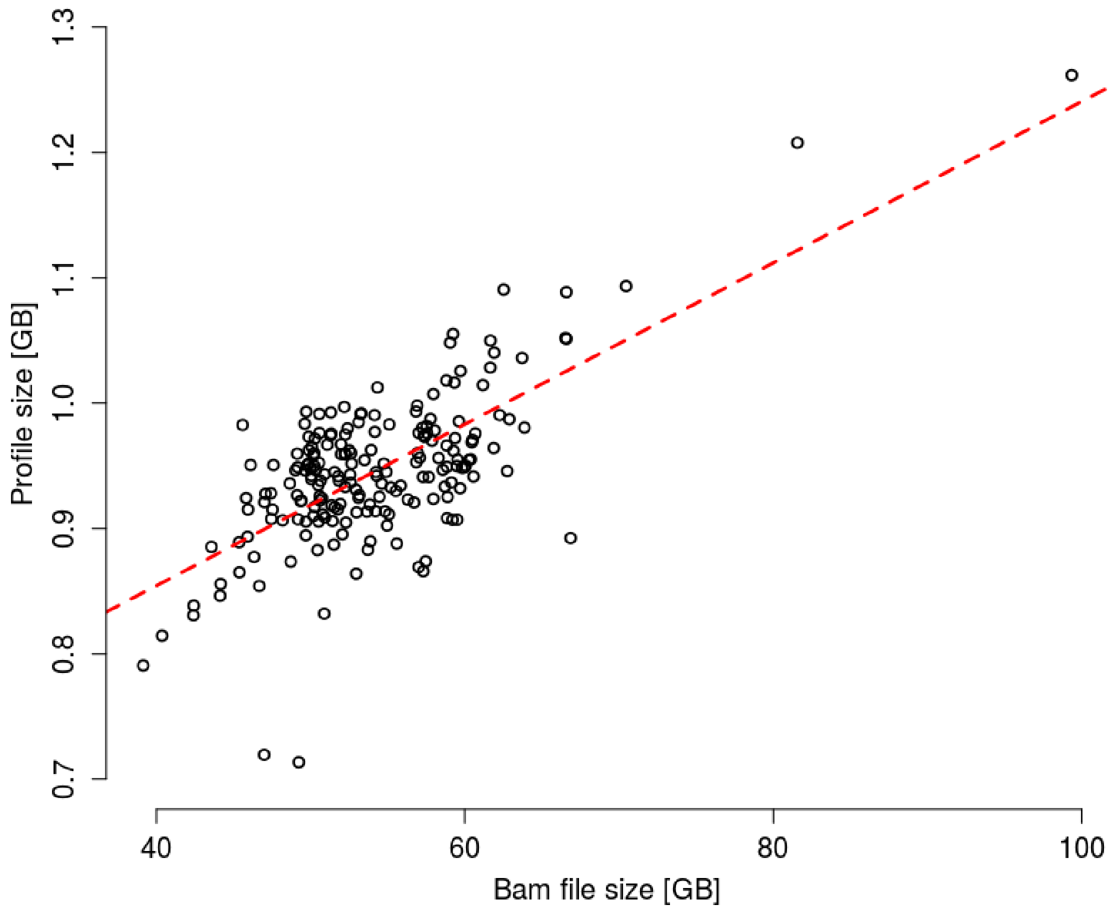

Supplementary figure 2 - Disk space used by PopDel's profile files compared to the disk space used by the original BAM files of all Polaris (Diversity and Kids cohort) samples. The red line indicates a linear regression line. The median profile size is 0.9 GB, which is 1.78% of the median BAM file size of 53.1 GB. The mean relative size of the profiles is 1.76%.

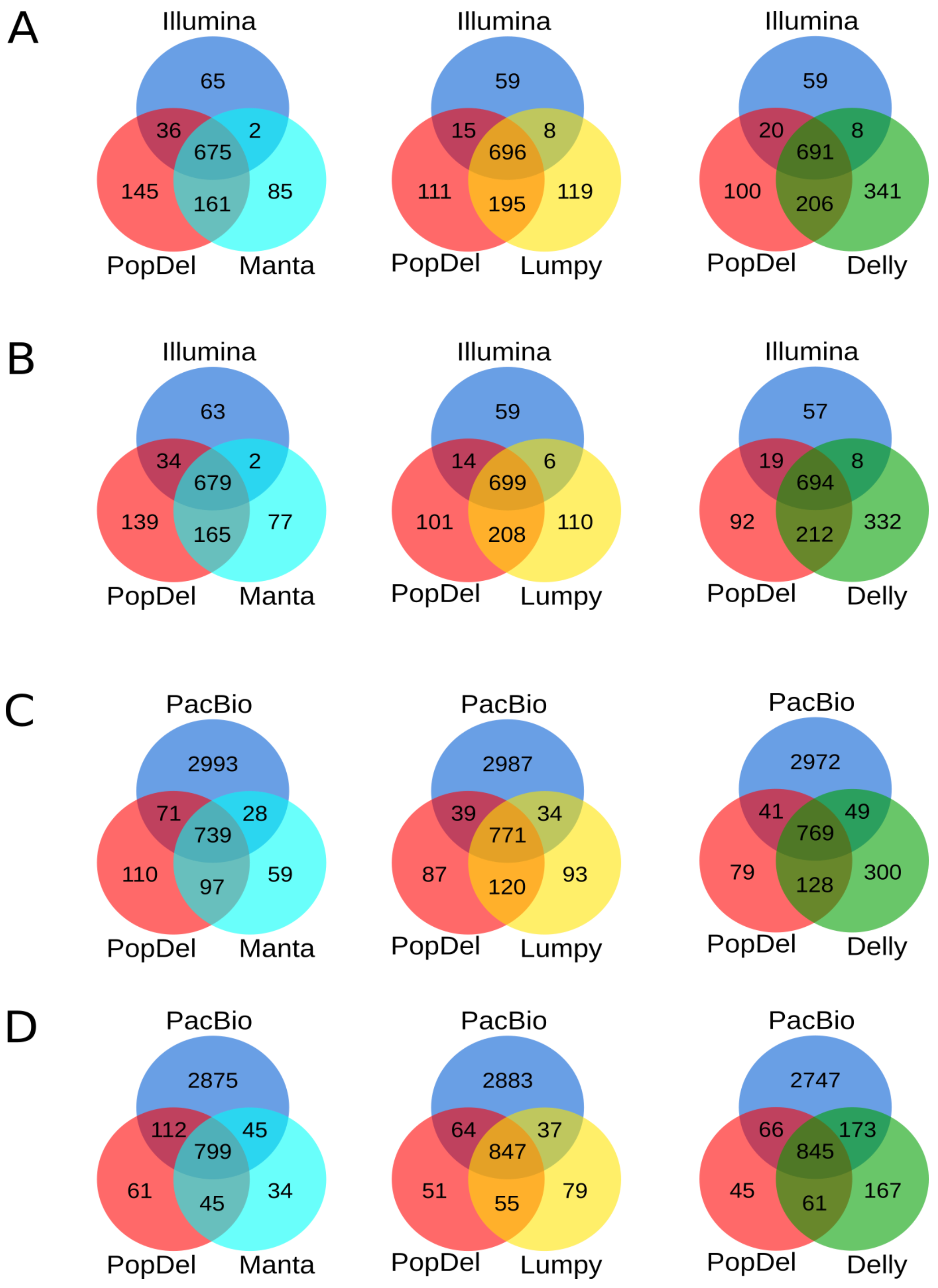

Supplementary figure 3 - Venn diagrams of call set overlaps on NA12878. Comparison with Illumina short read reference set (A, B) and PacBio long read reference set (C, D) for minimum reciprocal overlap of 80% (A, C) and 0.1% (B, D).

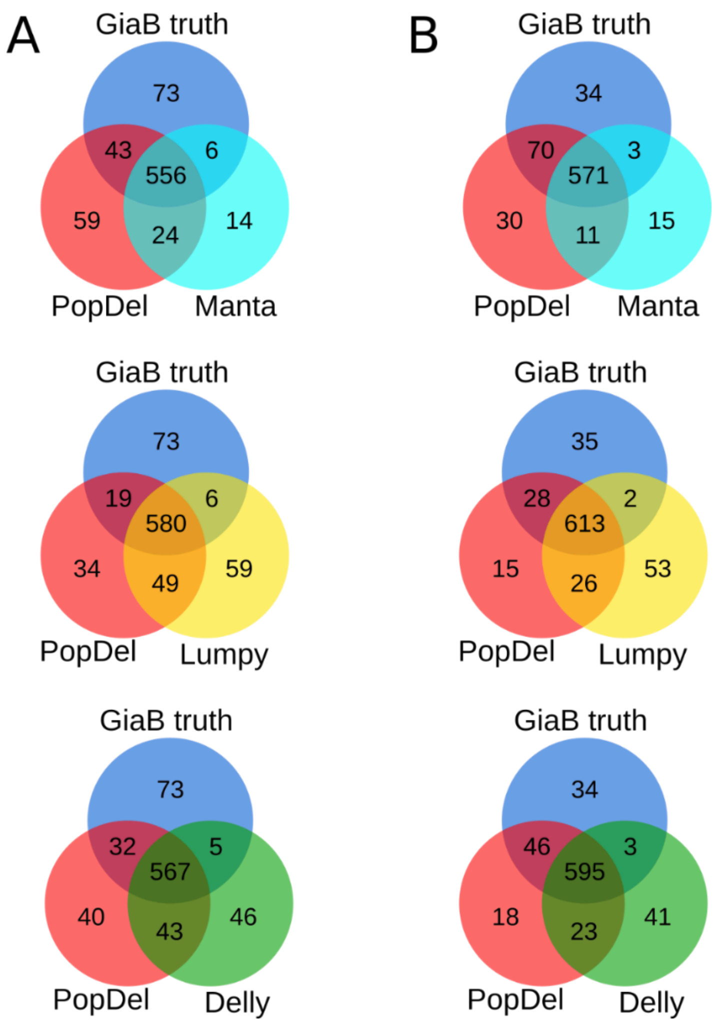

Supplementary figure 4 - Venn diagrams of call set overlaps on HG002. Comparison with GiaB truth set in high confidence regions for minimum reciprocal overlap of 80% (A) and 0.1% (B).

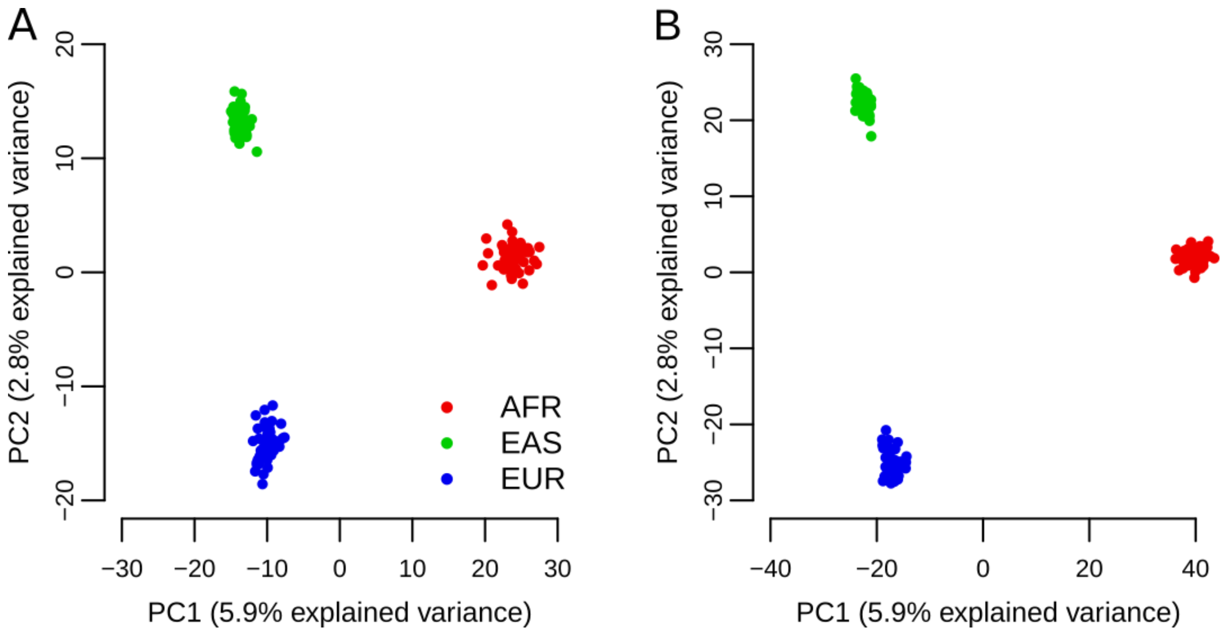

Supplementary figure 5 – PCA biplot for Delly (A) and Lumpy (B). PCA calculations were based on the variant allele counts of the tools for the Polaris Diversity cohort.

#### Supplementary text

### 1. Implementation details of PopDel

#### 1.1. Binary profile format

The binary format, defined in Table S1, stores insert sizes efficiently on disk. One important factor is avoiding the suboptimal representation of big integers as strings of chars.[1] Further disk space is saved by avoiding redundant information like the full name of the chromosome for each genomic position. Instead a 32-bit unsigned integer is used as identifier. Thanks to the strict size definition of every entry it is possible to avoid separators like whitespaces, commas, or colons otherwise necessary. This also enables efficient indexing and jumping in the file. Lastly, the stored insert sizes are grouped in genomic windows of 256 base pairs based on their start positions, as can be seen in Supplementary table 2. This way the positional information has to be stored only once per window and not per insert size. The offset of each read pair's position from the start position of the window is stored as a single char[2]. If one wants to inspect the profile, PopDel offers the function 'view' to convert the profile into a human readable format.

[1] One char typically takes 8 bits of disk space. If one wants to represent a number consisting of e.g. 7 digits, like it is to be expected when storing genomic coordinates, this adds up to $7 \cdot 8 = 56$ bits of space for this single number. By using a 32-bit unsigned integer for storing instead, PopDel can store the same number in 24 less bits.

[2] One char can hold the numbers from 0 to 255, which is the reason why windows of 256 bp are applied.

Table S1 - Definition of PopDel's binary profile format.

| Field | Description | Type | Value |
|---|---|---|---|
| magic | PopDel magic string | char[7] | POPDEL\1 |
| l_region | Index region size | uint32_t | |
| n_file_offsets | # file offsets in the index (index size) | uint32_t | |
| List of file offsets (n=n_file_offsets) | | | |
| file_offset | File offset (index entry) | uint64_t | |
| n_rg | # read groups | | |
| List of read group names and insert size histograms (n=n_rg) | | | |
| l_rg_name | Length of the read group name plus 1 (including NUL) | uint32_t | |
| rg_name | Read group name; NUL-terminated | char[l_rg_name] | |
| median_isize | Median insert size in this read group | int16_t | |
| stddev_isize | Standard deviation of insert size in this read group | int16_t | |
| read_length | Length of reads in this read group | uint16_t | |
| hist_start | First insert size listed in histogram | uint16_t | |
| hist_end | Last insert size listed in histogram plus 1 | uint16_t | |
| List of read pair counts per insert size (n=hist_end - hist_start); insert size histogram | | | |
| count | # read pairs with corresponding insert size | uint32_t | |
| n_ref | # reference sequences | uint32_t | |
| List of contig names (n=n_ref) | | | |
| l_ref_name | Length of the reference name plus 1 (including NUL) | uint32_t | |
| ref_name | Reference sequence name; NUL-terminated | char[l_ref_name] | |
| l_ref | Length of the reference sequence | uint32_t | |
| List of windows (until the end of the file) | | | |
| refID | Reference sequence ID, 0 ≤ refID < n_ref | uint32_t | |
| pos | Start position of the window (0-based) | uint32_t | |
| List of read group entries for the window (n=n_rg) | | | |
| n_rp | # read pairs | uint32_t | |
| List of read pairs for the read group in the window (n=n_rp) | | | |
| pos_offset | Offset of read pair position from window start position | Char | |
| isize_dev | Deviation from median insert size | int16_t | |

The first column describes the name of the field, whose content is described in the second column. Column 3 indicates the datatype and the size of the entry. Except for the magic string, which is used for identifying the profile format and version, no field has default value as indicated by column 4. A line saying "List of …(n=n_x)" indicates that the following block is repeated n times, potentially with different content.

#### 1.2. Buffering and length limitations during profile creation

PopDel creates the profile by checking the reads of the BAM file one after the other. If a read is in the forward orientation and its corresponding reverse read maps at most $l_{max} + \mu_S$ base pairs downstream (with $l_{max}$ as the maximum deletion length which defaults to 10,000 bp and $\mu_S$ as the median of the sample's insert sizes), the read is added to a buffer. If a read is in reverse orientation the corresponding forward read is removed from the buffer and the pair's insert size is added to the profile. This buffering ensures that both reads fulfil the requirements listed in Supplementary table 1 without jumping in the BAM file. The default value of 10,000 bp for $l_{max}$ has been selected to keep the buffering to a minimum and because we argue that deletions above a size of 10,000 bp can be easily found using read depth methods. This value can be increased by the user.

#### 1.3. Sampling regions for creating the insert size histogram

The sampling uses a set of 22 default regions, one 1,000,000 bp region from each autosome. The regions can also be specified by the user. PopDel parses all reads passing the filters in the first of the desired regions and counts the number of read pairs per insert size to obtain an empirical histogram of the insert sizes. If the minimum number of sampled read pairs (default 50,000 per read group) has been reached, no further regions are used for sampling. Otherwise the sampling continues with the next regions. The regions are listed in PopDel's wiki on GitHub.

#### 1.4. Robust estimation of median and standard deviation of insert size histogram

The median $\mu_S$ and standard deviation $\sigma_S$ of the insert size distribution are estimated and refined by trimming the histogram to the insert size interval $[\mu_S - 3\sigma_S, \mu_S + 3\sigma_S]$, then repeating the calculation and trimming again until both $\mu_S$ and $\sigma_S$ converge. The final histogram is called $H_{abs}^S$. The refinement mitigates the effect outliers when calculating $\mu_S$ and $\sigma_S$, contributing to the overall robustness of the estimation.

#### 1.5. Note on representations of numeric values during the calling

Due to the potentially large amount of data in a single window and the numerous multiplications of very small likelihoods, it is to be expected that numbers might occasionally drop below the smallest possible values for the data type double. To avoid this, the values of the histograms are stored as their log-values and all likelihood ratios are internally computed as log-likelihood-ratios. This results in an addition of the log-likelihood ratios instead of multiplications and a bigger domain for the possible values. The formulas in the Methods and Supplementary text nevertheless avoid the logarithms to keep the underlying formulas as concise as possible.

#### 1.6. Handling of multiple read groups in a sample

For some experiments, a sample might consist of multiple read groups, which further might originate from different libraries with distinct insert size distributions. PopDel handles this by creating one insert size histogram per read group instead of per sample. During the calling, PopDel handles all reads according to the read group they originate from, i.e. it compares the insert sizes to the corresponding distribution. All insert size histogram related parameters $(H(x), \mu, \sigma)$ as well as the reference shift $\epsilon$ and the weights $a_g$ are calculated and handled on a per read group basis.

For the calculation of the likelihoods we obtain:

$$\mathcal{L}(\text{no del}) = \pi \prod_{S\in\mathcal{S}} \prod_{\mathrm{R}\in\mathcal{R}^S} \mathcal{L}\left(G_0 | \Delta^{\mathrm{R}}\right)$$

$$\mathcal{L}(\text{del of length } l) = (1-\pi) \prod_{S\in\mathcal{S}} \prod_{\mathrm{R}\in\mathcal{R}^S} a_0^R \cdot \mathcal{L}\left(G_0 | \Delta^{\mathrm{R}}\right) + a_1^R \cdot \mathcal{L}\left(G_1 | \Delta^{\mathrm{R}}\right) + a_2^R \cdot \mathcal{L}\left(G_2 | \Delta^{\mathrm{R}}\right)$$

with $\mathcal{R}^S$ as the set of read groups of sample S, $\Delta^{\mathrm{R}}$ is the set of insert size deviations of read group $R$ in the current window and $\pi$ as the prior (default $10^{-4}$). We further have:

$$\mathcal{L}\left(G_0 | \Delta^{\mathrm{R}}\right) = \prod_{\delta\in\Delta^R} H^R(\delta - \epsilon_{\mathrm{R}})$$

$$\mathcal{L}\left(G_1 | \Delta^{\mathrm{R}}\right) = \prod_{\delta\in\Delta^R} \frac{H^R(\delta - \epsilon_{\mathrm{R}}) + H^R(\delta - \mathrm{l})}{2}$$

$$\mathcal{L}\left(G_2 | \Delta^{\mathrm{R}}\right) = \prod_{\delta\in\Delta^R} H^R(\delta - l)$$

where $H^R(x)$ denotes the transformed insert size histogram of read group $R$ (see section 1.8). The sample and genotype specific weights become read group and genotype specific weights:

$$a_g^R = \frac{\mathcal{L}\left(G_g | \Delta^{\mathrm{R}}\right) \cdot F\left(f, G_g\right)}{\sum_{j=0}^{2}\left(\mathcal{L}\left(G_j | \Delta^{\mathrm{R}}\right) \cdot F\left(f, G_j\right)\right)}$$

For the allele frequency $f$ we obtain:

$$f^{new} = \frac{1}{2|\mathcal{S}|} \cdot \sum_{S\in\mathcal{S}} \sum_{R\in\mathcal{R}^S} (a_1^R + 2a_2^R)$$

And we update the deletion length estimate as:

$$l^{new} = \frac{\sum_{S\in\mathcal{S}} \sum_{R\in\boldsymbol{\mathcal{R}}^S} \sum_{\boldsymbol{\delta}\in\Delta^R} \delta \cdot P_{l,\epsilon_R}^R(\delta)}{\sum_{S\in\mathcal{S}} \sum_{R\in\boldsymbol{\mathcal{R}}^S} \sum_{\delta\in\Delta^R} P_{l,\epsilon_R}^R(\delta)}$$

with

$$P^R_{l,\epsilon_R}(\delta) = a^R_1 \cdot \frac{H^R(\delta - l)}{H^R(\delta - \epsilon_R) + H^R(\delta - l)} + a^R_2$$

#### 1.7. Initialization of the deletion length and allele frequency

Given the insert size deviations of read pairs overlapping some genomic window, PopDel first suggests one initial deletion length per sample and then clusters these values across all samples into a small set of initial deletion lengths. More specifically, PopDel inspects the insert size deviations of each sample $S$ separately and selects the third quartile $Q_3(\Delta^S)$ as an initial length suggestion. If $Q_3(\Delta^S) \geq 4\sigma_S$, it is added to a list. After processing all samples, the values on this list are clustered such that values that differ by at most 50 bp end up in the same cluster. The means of all clusters are subsequently used as initial deletion lengths $l^{init}$. For each $l^{init}$, an initial allele frequency is calculated as:

$$f^{init} = \frac{\sum_{S\in\mathcal{S}} \sum_{\boldsymbol{\delta}\in\Delta^S} b_{l,2\sigma_S}(\delta)}{2|\mathcal{S}|}, \quad b_{l,2\sigma_S}(\delta) = \begin{cases} 1 & \text{if } x \geq \max\left(l - 2\sigma_S, \frac{l}{2}\right) \wedge \delta \leq l + 2\sigma_S \\ 0 & \text{else} \end{cases}$$

where $|\mathcal{S}|$ denotes the total number of samples.

#### 1.8. Transformation of insert size histogram

The histogram $H^S_{abs}$, which has been created during the profiling step, stores absolute counts of observed insert sizes for sample $S$. Let $H^S_{abs}(i)$ return the number of counts for an insert size $i$. We compute a transformed histogram:

$$H^S_{trans}(\delta) \coloneqq H^S_{abs}(\delta + \mu_S) \cdot \frac{w + \max(\delta + \mu_S, 0)}{w}$$

Since PopDel subsequently only uses the deviation of insert sizes from the mean and not the insert sizes themselves, $H^S_{trans}$ takes the deviation $\delta$ of the insert size as an argument. The probability density function for the distribution is then calculated as follows:

$$H^S(\delta) \coloneqq \frac{H^S_{trans}(\delta)}{\sum_j H^S_{trans}(j)}$$

In the cases where the sampled distribution would return 0, a pseudo-count of default $\frac{\max(H^S)}{500}$ is applied instead. This is necessary to avoid a division by 0 or trying to take the logarithm of 0 in subsequent steps.

The reasoning behind this transformation is as follows. Since a bigger insert size and a higher window size increases the frequency of a read pair overlapping a window, we scale the values returned by $H_{abs}^{S}(i)$ with respect to insert size and window size. The number of possible mapping locations so that a read pair with insert size $i$ and read length $r$ overlaps the window of length $w$ is $w + \max(i - 2r, 0)$. This is because PopDel considers a read pair to overlap a window, if any part between the right end of the forward read and the left end of the reverse read maps to the window. The reads themselves are excluded. In case the forward and reverse read overlap, PopDel takes the rightmost alignment position of the forward read as a reference point. By dividing the size of the overlap by the window length, we gain the average number of consecutive windows, a read pair with insert size $i$ overlaps.

#### 1.9. Reference shift

Local biases such as GC-content can result in a local shift of the insert size distribution. To avoid that those shifts result in a false positive deletion call of the window, we introduced the reference shift $\epsilon$. To this end we allow the subtraction of $\epsilon$ from $\mu_S$ in the calculation of the likelihood for the reference model. $\epsilon$ is updated in a similar fashion as the deletion length, however without the sum across all samples, because $\epsilon$ is specific for each individual sample:

$$\epsilon_S^{new} = \frac{\sum_{\delta \in \Delta^S} \delta \cdot P_{d,\epsilon_S}^{S}(\delta)}{\sum_{\delta \in \Delta^S} P_{d,\epsilon_S}^{S}(\delta)}$$

As we expect only small biases, we limit the maximum value for $\epsilon$ to the standard deviation $\sigma_S$ of the sample's insert size distribution. Larger values may originate from homozygous deletions and interfere with the size estimations. Therefore, if $\epsilon$ becomes greater than $\sigma_S$, it is set to $0$. Setting $\epsilon$ to $\sigma_S$ instead, can distort the predicted length of homozygous deletions by this value.

#### 1.10. Termination of iterative parameter estimation

The iteration of PopDel's window-wise calling stops if the new estimates for the pair of deletion length and frequency has been observed before, the maximum number of iterations (default 15) is exceeded, the allele frequency drops below a threshold (default $10^{-10}$) or the estimate for the deletion length drops below a minimum length (default 95% quantile of 4 times all standard deviations of all samples' insert size histograms). The two latter cases result in no call being made for the respective window for that specific deletion. In case two pairs of estimates for frequency and deletion length alternate during the iteration, PopDel assumes the pair yielding the higher likelihood ratio to be the correct one.

#### 1.11. Deletion start position estimation

During the iteration in one window, PopDel keeps track of the positions of read pairs whose insert size deviation from the sample mean supports the deletion estimate. Let $Q_{99}(H^S)$ be the 99% quantile of the distribution of insert size deviations of sample $S$. A read pair overlapping the window with insert size deviation $\delta$ is considered to support a deletion of length $l$ if $l - Q_{99}(H^S) \leq \delta \leq l + Q_{99}(H^S)$. The positions of deletion supporting read pairs are stored in a sorted list and the 80% quantile of this list is taken as the start position of the variant. This provides us with a precise estimate of the deletion start position, since the position of the read pairs stored by PopDel is the rightmost aligned position of the forward read in each pair. Therefore, PopDel implicitly considers the split-reads for estimating the precise start position.

We used our simulated data to quantify the exactness of this approach. We calculated the deviation of the tools' predicted start positions from the actual simulated position in the first 10 samples of the random deletion simulation (Methods, Supplementary text section 2.1.7). Figure S1 shows that our approach is on par with the other tools that explicitly consider split-read alignments.

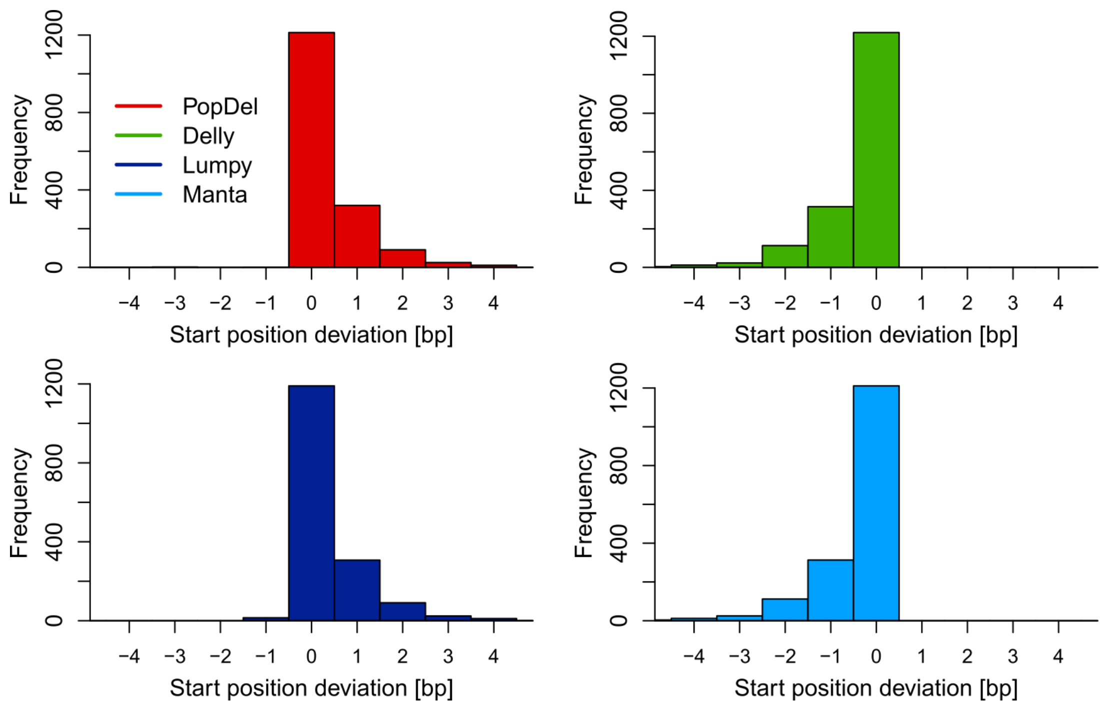

Figure S1 – Histograms of deletion start position deviations. The deviation was calculated as the difference between predicted and simulated position of the variant. A negative deviation indicates that the predicted position is upstream of the simulated position. Variants were not left or right aligned.

#### 1.12. Conditions for the combination of consecutive windows

Let $w_i$ be the first of $n$ windows for which the null hypothesis can be rejected and let $w_{i+k}, 0 < k < n$ be another such window. Let $l_i$ and $l_{i+k}$ be the deletion length estimates for $w_i$ and $w_{i+k}$ respectively. Let $r_i := (\text{delStart}(w_i), \text{delEnd}(w_i))$ denote the range spanned by the leftmost and rightmost alignment positions of the read pairs whose insert sizes support $w_i$ and let $r_{i+k}$ be defined analogously. When considering one window after the other, we test if the following conditions for $w_i$ and $w_{i+k}$ are met:

a) $|\text{l}_\text{i} - l_{i+k}| \leq \max\left(\frac{\min(l_i, l_{i+k})}{2}, 2\bar{\sigma}\right)$, with $\bar{\sigma}$ as the mean of all $\sigma_S$ of all samples.

b) $r_i$ and $r_{i+k}$ overlap by at least $\min\left(\frac{\min(r_\text{i}, r_{i+k})}{4}, \min(r_\text{i}, r_{i+k}) - 2\bar{\sigma}\right)$.

If both a) and b) are satisfied, $w_i$ and $w_{i+k}$ are combined. If only a) is satisfied, an additional check is preformed to see if the deletion is interrupted:

c) $(|r_i| < l_i \vee |r_{i+k}| < l_{i+k}) \wedge \text{delStart}(w_{i+k}) - \text{delStart}(w_i) < \min(l_i, l_{i+k}) + 4\sigma$

where $|r_i|$ denotes the length of the spanned range. If a) and c) are true, the deletion end position in the combined window is updated to match that of $w_{i+k}$.

#### 1.13. Additional filters during joint calling

PopDel applies additional filters on the final variant calls to improve the precision of the results. The first filter is applied during the iterative parameter estimation. If in a window a sample (over all read groups) has a coverage below two, no deletion will be initialized from the data of this sample and PopDel will not attempt to genotype the sample later on in the respective window. We assume that this window is unreliable if more than 90% of the samples in the window are lacking coverage and filter the window accordingly. In the case of too high coverage (default at least 100 read pairs of a single read group overlapping one window) the read pairs of this read group are ignored in all calculations. Samples consisting solely of read groups with high coverage in one window will also not be assigned genotypes for the respective window. Another filter, the *relative window coverage,* is the value obtained by dividing the product of window length and number of combined windows $M$ by the estimated deletion length. By default, PopDel requires

$$\frac{\text{M} \cdot w}{l} \geq 0.5$$

Lastly, the variant is filtered if all samples are genotyped as non-carriers.

#### 2. Evaluation on simulated and real data

To assess the performance of PopDel, Delly, Lumpy, Manta and GRIDSS the tools were compared on the different simulated and real data sets described in the Methods. A more detailed description of the setup of the callers and the evaluation is described in the following sections.

##### 2.1. Simulated data with random deletions

The simulation of random variants was performed as described in the Methods. Scripts for the reproduction of the workflow can be found in the GitHub repository (see URL's of the Main text). The setup of the callers is described in the following sections.

###### 2.1.1. Running PopDel on simulated data with random deletions

The PopDel workflow for all tests consisted of the following steps

1. Apply `popdel profile` on each sample individually
2. Apply `popdel call` jointly on all profiles

The option `–r chr21` for limiting the processing to chromosome 21 was applied. No further filtering of the calls was performed.

###### 2.1.2. Running Delly on simulated data with random deletions

Delly was applied jointly on all simulated samples using default parameters, except for the `-n` option for disabling the small-indel realignment. All non-deletion variants were removed prior to evaluation. No filtering for "QUAL==PASS" was applied, because doing so had a negative impact on the performance of Delly on the simulated data.

###### 2.1.3. Running Lumpy on simulated data with random deletions

Lumpy was applied via Smoove, as recommended by the authors. The recommended file for excluding regions on GRCh37/GRCh38 was used. Additionally, all mappings to chromosomes other than chromosome 21 were ignored in the random deletion simulation by using the expression
`'chr1,chr2,chr3,chr4,chr5,chr6,chr7,chr8,chr9,chr10,chr11,chr12,chr13,chr14,chr15,chr16,chr17,chr18,chr19,chr20,chr22,chrX,chrY,chrM,chrEBV,~.*_.*'`
with the option `--excludeChroms`. The Smoove pipeline consisted of the following steps:

1. Single sample calling and genotyping with `smoove call` for each sample individually (`--processes 1`)
2. Merging the calls of all samples with `smoove merge`

3. Genotyping of each individual sample against the merged calls with `smoove genotype`.
4. Joining the calls of all samples using `smoove paste`.

All non-deletion variants were removed prior to evaluation. No further filtering was performed.

##### 2.1.4. Running Manta on simulated data with random deletions

Manta's workflow consisted of two steps:

1. Configuration via `configManta.py` for all bam files together (`--region chr21`)
2. Joint calling via `runWorklow.py` with `-m local –j 1` (local machine and one thread)

All non-deletion variants were removed prior to evaluation. No filtering for "QUAL==PASS" was performed, because doing so had a negative impact on the performance of Manta on the simulated data. Because the additional variants in the "candidate" file had a very high false positive rate, they were not added to the final output.

##### 2.1.5. Running GRIDSS on simulated data with random deletions

GRIDSS was applied on all samples jointly with an 8 GB heap size for the JVM and one thread. Variants with "FILTER==LOW_QUAL" were removed. An adapted version of the annotation script provided in GRIDSS' GitHub repository[3] was used to convert the break points into deletion calls.

##### 2.1.6. Evaluation on simulated data with random deletions

A variant called by one of the tools was considered to be a true positive (TP) if `bedtools intersect` reported a reciprocal overlap of least 50% between the area spanned by the called variant and the area spanned by some true simulated variant. All simulated variants in the truth set that hat no such overlap were counted as false negatives (FN). If no such overlap existed, it was considered to be a false positive (FP). Each simulated variant was only allowed to be matched with one predicted variant. Genotypes of different samples were not considered in this evaluation.

##### 2.1.7. Alternative evaluation with genotype consideration

In addition to the above approach we applied an alternative evaluation metric that also takes the genotypes of the different samples into account. A called variant was considered to match a simulated true variant if the start position of the call was within 300 bp of the simulated variant's start position and the predicted length of the call was within 150 bp of the real deletion

[3] https://github.com/PapenfussLab/gridss/blob/master/example/simple-event-annotation.R

length. Further, the genotypes were inspected. If the predicted genotype of a sample was equal to the simulated genotype, it was counted as two correct alleles. If the predicted genotype did not match the simulated genotype and both genotypes were homozygous, two incorrect alleles were counted. If the predicted genotype did not match the simulated genotype and one of the genotypes was heterozygous, one correct and one incorrect allele was counted. If a simulated variant had no matching predicted variant, incorrect alleles were counted accordingly. Predicted 0/0 genotypes were not counted as correct alleles, because otherwise a tool could arbitrarily boost its score by calling a "variant" that was not present in the truth set and giving all 0/0 genotypes. The resulting plots of this approach are displayed in Figure S2. Because GRIDSS does not produce genotypes, it was excluded from the genotype evaluation.

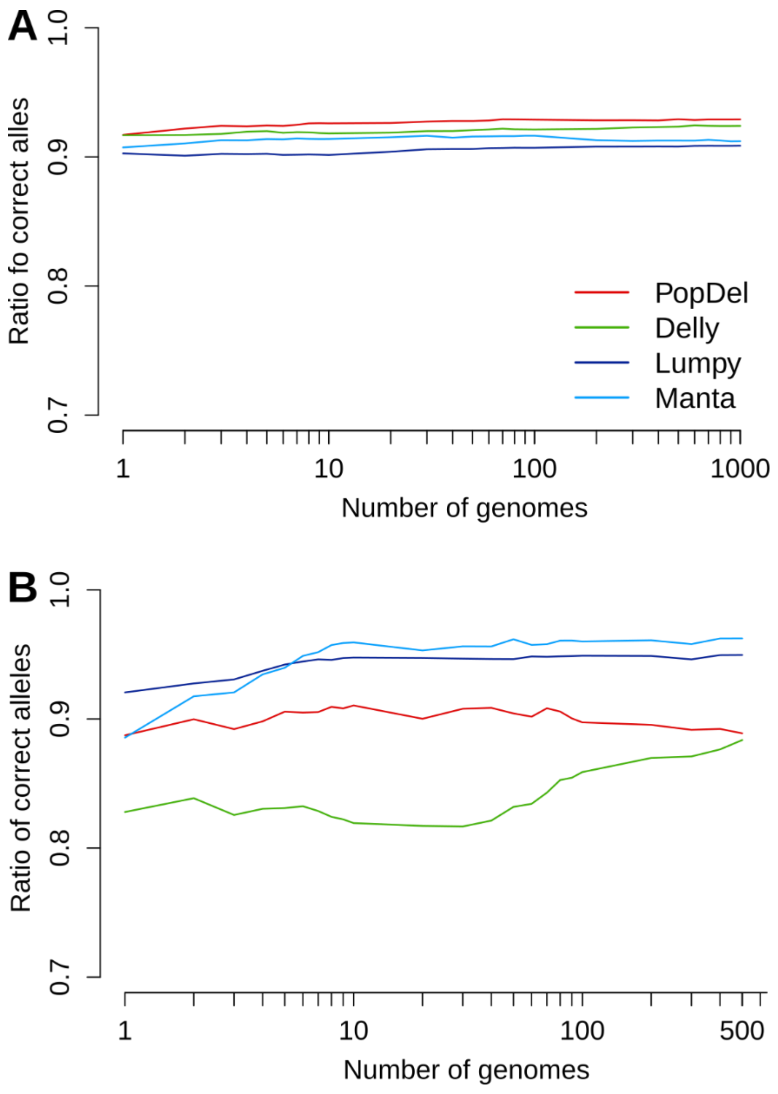

Figure S2 – Genotyping evaluation based on alternative overlap evaluation approach (max pos-delta=300, max length-delta=150) on random chromosome 21 deletions (A) and G1k deletions inserted into simulated chromosomes 17 to 22 (B). All genotyped alleles were counted and compared to the simulated haplotypes. Any incorrect allele was counted as an error.

##### 2.1.8. Assessment of running time and memory consumption

Resource consumption of all tools for all tests on simulated data was measured using `/usr/bin/time`. CPU time was calculated as the sum of CPU seconds the process spent in user mode (%U) and the number of CPU seconds the process spent in kernel mode (%S). Memory consumption was measured as maximum resident set size of the process during its lifetime (%M). Wall clock time was measure as the elapsed time (%e).

##### 2.1.9. Tradeoff between running time and memory consumption for PopDel

PopDel offers the option for individually setting the maximum number of buffered windows. This allows to manage the memory consumption of PopDel while also influencing the running time in a direct manner. For measuring the effect of varying buffer sizes on running time and memory consumption, PopDel was applied with varying settings for the buffer size on 100 samples simulated from chromosome 21 with random deletions. The results are displayed in Figure S3 below.

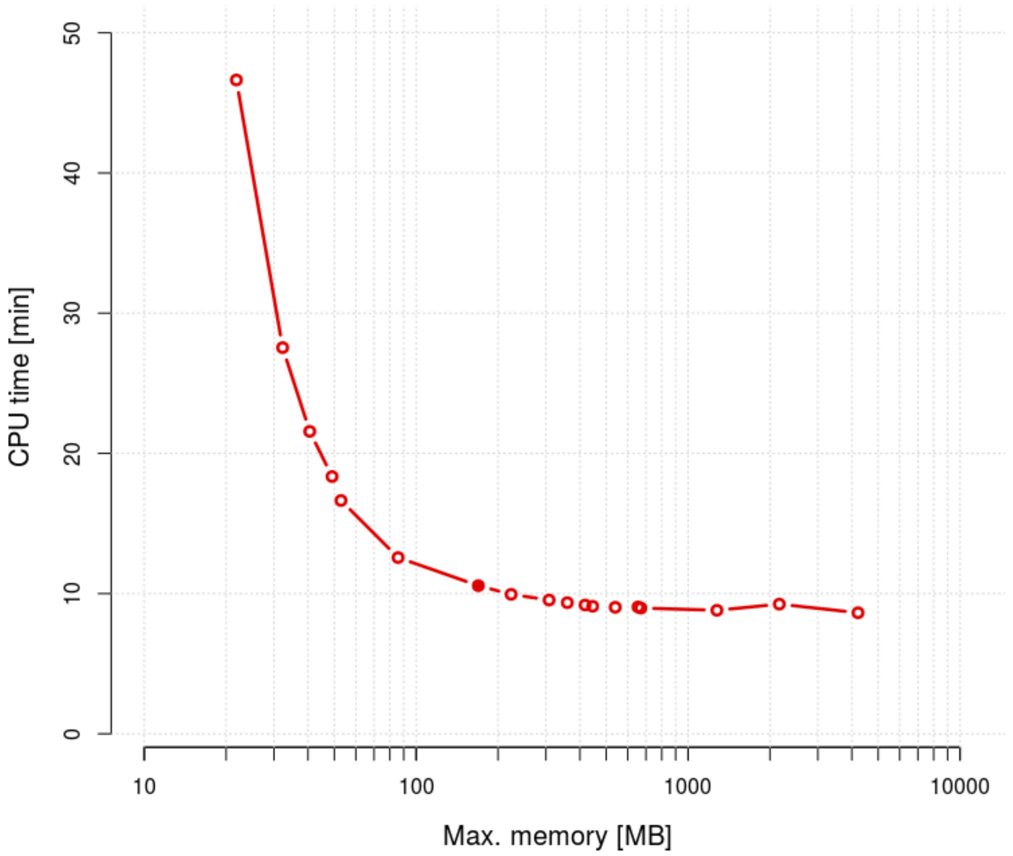

Figure S3 – PopDel call memory consumption compared to its CPU time for varying buffer sizes on 100 samples simulated from chromosome 21 with random deletions. The filled red dot indicates the default option for the buffer size (200,000 windows).

#### 2.2. Simulated 1000 Genomes Project data

The simulation of variants taken from the 1000 Genomes Project was performed as described in the Methods. Scripts for the reproduction of the workflow can be found in the GitHub repository (see URL's of the Main text). The setup of the callers was the same as described in sections 2.1.1-2.1.5, with the exception that chromosomes 17 to 22 were considered. The same evaluation criteria as described in 2.1.6 were applied.

#### 2.3. Evaluation of single and trio wise calling using NA12878 and the HG002 trio

In a first comparison of the performance on real data, we evaluated the precision and recall and the call set overlaps between the tools' predictions and reference deletion sets for NA12878, HG002. The accession numbers, procedure of the mapping, setup of the callers and evaluation metrics are described in the following sections.

##### 2.3.1. Accession numbers for HG002 trio data

Because the HG002 trio was sequenced to a very deep coverage, we decided to use only read data from the first three flow cells of the GiaB data repository. The resulting alignments for HG002, HG003 and HG004 reached an average coverage of 45x, 68x and 72x, respectively. The accession numbers of the FASTQ files are as follows:

HG002:

SRR1766443, SRR1766444, SRR1766445, SRR1766455, SRR1766456, SRR1766457, SRR1766467, SRR1766468, SRR1766469, SRR1766479, SRR1766480, SRR1766481, SRR1766491, SRR1766492, SRR1766493, SRR1766503, SRR1766504, SRR1766505, SRR1766515, SRR1766516, SRR1766517, SRR1766527, SRR1766528, SRR1766529, SRR1766539, SRR1766540, SRR1766541, SRR1766555, SRR1766556, SRR1766557, SRR1766566, SRR1766567, SRR1766568, SRR1766582, SRR1766583, SRR1766584.

HG003:

SRR1766542, SRR1766543, SRR1766545, SRR1766546, SRR1766576, SRR1766577, SRR1766579, SRR1766580, SRR1766589, SRR1766590, SRR1766592, SRR1766593, SRR1766596, SRR1766597, SRR1766599, SRR1766601, SRR1766602, SRR1766603, SRR1766615, SRR1766627, SRR1766639, SRR1766651, SRR1766663, SRR1766675, SRR1766687, SRR1766699, SRR1766700, SRR1766701, SRR1766713, SRR1766714, SRR1766726, SRR1766727, SRR1766728, SRR1766740, SRR1766741, SRR1766742

HG004:

SRR1766763, SRR1766764, SRR1766765, SRR1766775, SRR1766776, SRR1766777, SRR1766787, SRR1766788, SRR1766789, SRR1766799, SRR1766800, SRR1766801, SRR1766811, SRR1766812, SRR1766813, SRR1766823, SRR1766824, SRR1766825, SRR1766835, SRR1766836, SRR1766837, SRR1766847, SRR1766848, SRR1766849, SRR1766850, SRR1766860, SRR1766861, SRR1766871, SRR1766872, SRR1766873, SRR1766883, SRR1766884, SRR1766885, SRR1766895, SRR1766896, SRR1766897

##### 2.3.2. Mapping of real data

Reads of NA12878 were mapped to GRCh38 and reads of HG002 and his parents to GRCh37 using `bwa mem` with its '`-M`' and '`-R`' options. SAM to BAM conversion, sorting and indexing was performed using Samtools. Duplicates were marked using Picard tools.

##### 2.3.3. Running PopDel on NA12878 and the HG002 trio

Creation of the profiles was performed using the default options of `popdel profile`. For NA12878, calling was performed in parallel on chromosomes 1 to 22 and the X chromosome using the option '`-r`'. For the HG002 trio, calling was performed in parallel on chromosomes 1 to 22 using the option '`-r`'. As the HG002 trio has a quite high coverage, the maximum coverage for PopDel to consider was set to the three-fold mean coverage of each sample using the option '`-A`'.

Example command line:

```
popdel profile sample.bam;
popdel call profile_paths.txt -r chr1 -o sample.chr1.vcf;
```

##### 2.3.4. Running Delly on NA12878 and the HG002 trio

Calling was performed using Delly's recommended options. For NA12878, the recommended file of excluded regions on GRCh38 was used and only chromosomes 1 to 22 and the X chromosome were considered by adding the Y chromosome to the file of excluded regions. The HG002 trio was analyzed with the recommended file of excluded regions on GRCh37.

Example command line:

```
delly call -n -x human.hg38.excl.tsv -o sample.bcf -g ref.fasta sample.bam;
```

##### 2.3.5. Running Lumpy on NA12878 and the HG002 trio

Calling was performed using the recommended options of Smoove. For NA12878, the recommended file of excluded regions on GRCh38 was used and only mappings to chromosomes 1 to 22 and the X chromosome were considered. For the HG002 trio, the recommended file of excluded regions on GRCh37 was used and only mappings to chromosomes 1 to 22 were considered.

Example command line:

```
smoove call -x −−outdir sample.out −−name sample.name \
−−exclude exclude.cnvnator_100bp.GRCh38.20170403.bed −−excludeChroms \
‘chrY,chrM,chrEBV,~.*_.*’ −−fasta ref.fasta -p 1 --genotype sample.bam;
```

##### 2.3.6. Running Manta on NA12878 and the HG002 trio

Calling was performed using Manta’s recommended options. For NA12878 the calling was limited to chromosomes 1 to 22 and the X chromosome.

Example command line:

```
configManta.py −−bam sample.bam --referenceFasta ref.fa \
--runDir workingdir --callRegions regions.bed.gz;
workingdir/runWorkflow.py -m local -j 1;
```

##### 2.3.7. Evaluation using `bedtools intersect`

Called variants and the reference deletion sets were converted to BED format and filtered for the desired length range (500-10,000 bp). Any variants with any overlap with a centromeric region (determined by `bedtools intersect`) were removed. Subsequently, overlaps were generated in a hierarchical fashion as displayed in Figure S4..

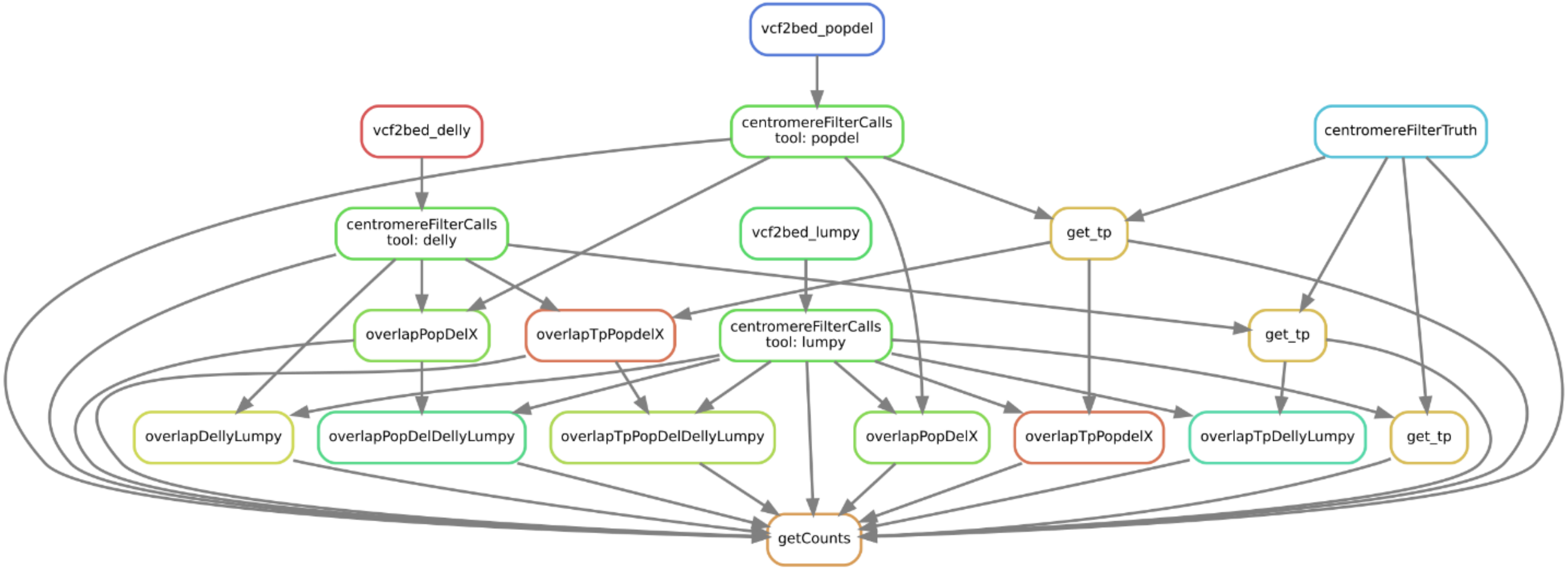

Figure S4 - Overview of the overlap generation workflow (created with Snakemake[4]).

---

[4] Köster, J., & Rahmann, S. (2012). Snakemake—a scalable bioinformatics workflow engine. *Bioinformatics*, *28*(19), 2520-2522.

##### 2.3.8. Analysis of the Mendelian inheritance error rate

Similar to the approach described for the Polaris Kids cohort in the Main text and below in section 2.4.5, we assessed the Mendelian inheritance error rate on the data of the HG002 trio when filtering for increasing genotype quality (GQ) thresholds (see Figure S5). Due to the low number of variants in this single trio we observe large fluctuation in the error rates

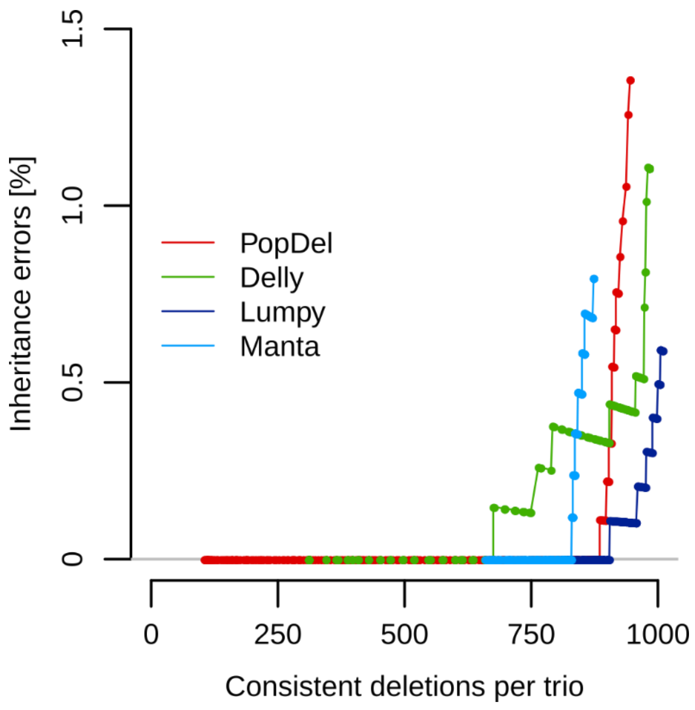

Figure S5 - Mendelian inheritance error rate by number of deletion sites in the HG002 trio that are consistent with Mendelian inheritance.

#### 2.4. Evaluation on the Polaris Diversity and Kids cohort

We applied PopDel, Delly and Lumpy on the 150 samples of the Polaris Diversity cohort and the 49 trios of the Polaris Kids cohort as described in the following sections. As we could not produce a workflow for Manta which finished on that many whole genomes, we do not provide a workflow here.

##### 2.4.1. Running PopDel on the Polaris Diversity and Kids cohort

The same default approach as described for NA12878 in section 2.3.3 was applied. Chromosomes 1 to 22 and the X chromosome were considered for the Diversity cohort and chromosomes 1 to 22 were considered for the Kids cohort.

##### 2.4.2. Running Delly on the Polaris Diversity and Kids cohort

As jointly calling all 150 samples of the Diversity cohort or 49 trios of the Kids cohort did not produce any results within 4 weeks (option '`-n`' enabled), a sample-wise approach had to be chosen for Delly. It was comprised of the following steps:

1. Sample-wise calling (`delly -n`)
2. Filter all non-deletion variants (`bcftools filter`)
3. Merge all variants of all samples (`delly merge`)
4. Sample-wise genotyping against the list of merged variants (`delly genotype`)
5. Filter variants with invalid positions (`bcftools filter`)
6. Sort and index filtered variants
7. Merge the genotyped calls of all samples (`bcftools merge`)
8. Index the BCF file of all genotyped variants
9. Apply germline filter (`delly filter`)

Steps 5 and 6 were necessary due to a known bug in Delly[5] causing some variants to be have POS==0. This made the resulting BCF files impossible to sort and index with BCFtools, which was necessary for merging and applying the germline filter.

##### 2.4.3. Running Lumpy on the Polaris Diversity and Kids cohort

The same default approach as for the simulated data sets (section 2.1.3) was applied, except that chromosomes 1 to 22 and the X chromosome were considered for the Diversity cohort and chromosomes 1 to 22 for the Kids cohort. Further, all non-deletion variants were removed from the call sets after single-sample calling with `smoove call`.

##### 2.4.4. Principal Component Analysis (PCA)

The variants were converted into a sample by variant matrix $M$. Field $M_{i,j} \in \{0,1,2\}$ contains the number of variant alleles sample $i$ carries for variant $j$. Duplicate rows were removed as they were uninformative for the PCA. Linkage disequilibrium (LD) was accounted for by calculating the correlation of neighboring variants and removing those with a significant Spearman correlation coefficient (P value threshold 0.05). The procedure was applied iteratively until no more significant correlations were found. From initially 12,203 variants, 4,339 were removed because they had the same allele counts as another variant for all samples (duplicate rows). 3 more variants were removed because they had the same counts for all samples. LD filtering removed 1,589 additional variants, leaving 6,272 variants (48.6%) for the

---

[5] https://github.com/dellytools/delly/issues/106

final PCA. PCA was computed using the R-function `prcomp` with the options for 0-centering and unit variance scaling of the variables enabled. PCA was also performed for the genotypes produced by Delly and Lumpy producing similar results (Supplementary figure 5).

##### 2.4.5. Genotype quality filtering for Mendelian error rate and transmission rate

For assessing the effect of filtering on the Mendelian inheritance error rate, we applied an increasing genotype quality (GQ) threshold. Further, all deletions that were not in Hardy-Weinberg equilibrium (P value threshold 0.01) were removed. All genotypes below the required GQ were excluded. Because most tools calculate the GQ in a different way, we determined at which point the Mendelian inheritance error rate of the tools dropped below 0.3% to calibrate the filter for the tools in a fair fashion. The GQ values for the tools were 26 (PopDel), 28 (Delly) and 78 (Lumpy). Those filter settings were also used during the calculation of the transmission rates (Table 1).

##### 2.4.6. Transmission rate pattern observed for Lumpy

The deletion genotypes predicted by Lumpy in the Polaris Kids cohort do not reach the expected transmission rate of 50% in our analysis (Figure 5) but stay below 48.3% independent of the genotype quality threshold we apply. A transmission rate below 50% can be explained by two scenarios:

1. False negative calls in the child when a heterozygous genotype was correctly called in one parent.
2. False positive calls in one of the parents when the child was correctly determined as a non-carrier.

We think that false positive calls in one of the parents are more likely than genotyping errors in the child when one parent is correctly called heterozygous.

#### 2.5. Evaluation on Icelandic data

We evaluated PopDel on different datasets of deCODE Genetics at several stages of PopDel's development.

##### 2.5.1. Transmission and Mendelian inheritance error on Icelandic data

PopDel v1.0.6 was applied on WGS data of 6,794 Icelandic trios. For all variants with a genotype quality threshold above 26, the Mendelian inheritance error rate and transmission rate were calculated considering only variants occurring in a single family with one parent being heterozygous for the variant and the other parent carrying only the reference allele.

##### 2.5.2. Sanity checks on Icelandic data

An early version of PopDel (V0.1-alpha-c74a6c0) was applied on WGS data of 4,959 Icelanders to examine the degree of agreement of PopDel's results with expectations based on previous studies and PopDel's behavior on big datasets.

A non-trivial task when performing variant calling on population scale data is the detection of overlapping variants, at potentially low allele frequencies. Figure S6 shows an example common deletion completely overlapping a small deletion with a low allele frequency (2 of 9,918 alleles). PopDel could identify and correctly genotype the two variants using joint calling.

The distribution of the number of variants per genome is expected to follow a normal distribution. The variants identified by PopDel follow this distribution as shown in Figure S7. The two visible peaks are thought to originate from sequencing technology related properties.

Further, more smaller deletions are expected to be present than bigger ones. Figure S8 shows that PopDel identifies exponentially more small deletions than bigger deletions.

A similar behavior is expected and can be observed for the allele frequency, which is shown in Figure S9.

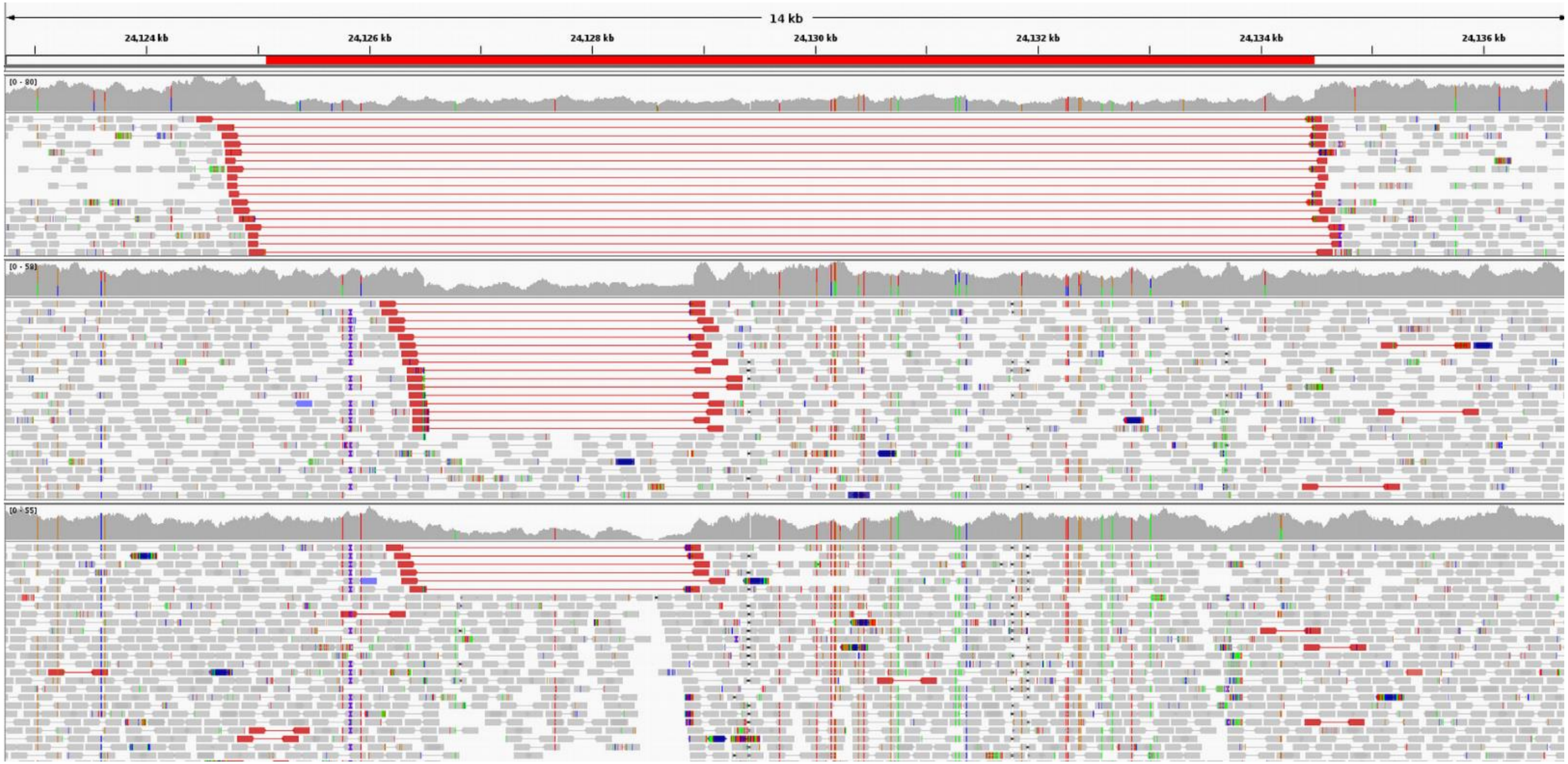

Figure S6 – Example of a rare deletion completely overlapped by a longer common deletion. The smaller deletion (middle and bottom) was only present in two of the analyzed 4,959 genomes, while the longer deletion (top) was present at a much higher allele frequency.

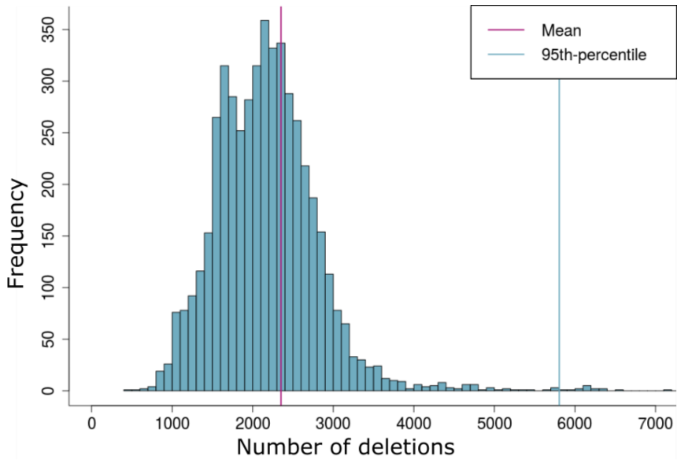

Figure S7 – Distribution of deletions per genome as identified jointly by PopDel on 4,959 Icelandic genomes.

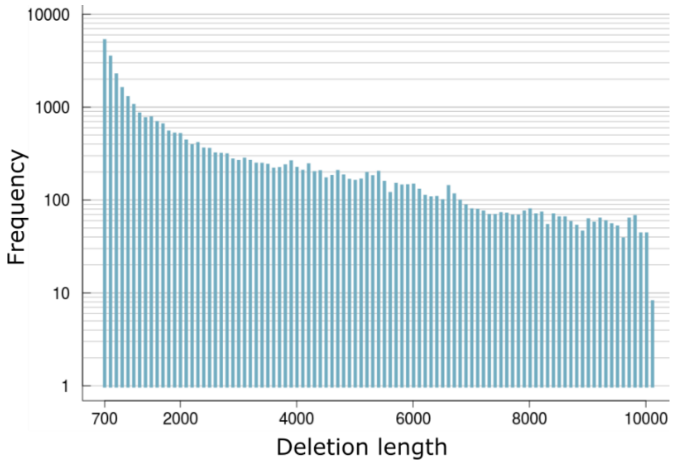

Figure S8 – Histogram of deletions lengths identified jointly by PopDel on 4,959 Icelandic genomes.

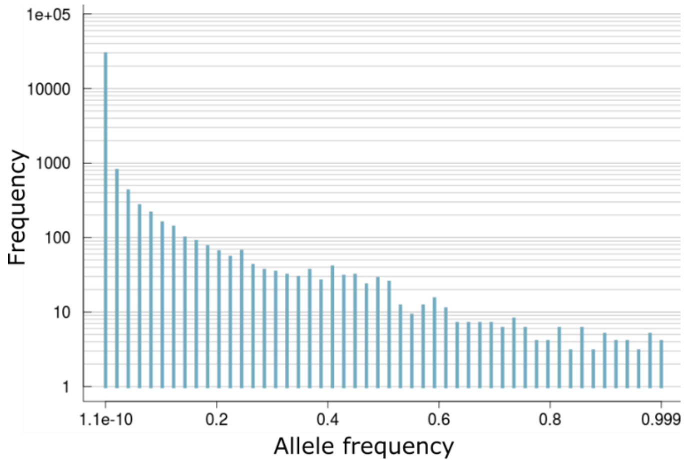

Figure S9 – Histogram of allele frequencies of deletions identified jointly by PopDel on 4,959 Icelandic genomes.
